# Microbial drama in four acts - Extreme rain events cause cyclic succession in plankton communities

**DOI:** 10.1101/2020.07.23.217935

**Authors:** Tanja Shabarova, Michaela M. Salcher, Petr Porcal, Petr Znachor, Jiří Nedoma, Hans-Peter Grossart, Jaromír Seďa, Josef Hejzlar, Karel Šimek

**Author notes:** corresponding author: Tanja Shabarova, mailing address: Biology Centre CAS, Institute of Hydrobiology, Na Sádkách 7, 370 05 České Budějovice, Czech Republic.

## Abstract

Highly abundant, small waterbodies contribute substantially to global freshwater shoreline and surface area. They are strongly interlinked with the terrestrial surrounding, thus controlling the flow of energy, nutrients and organisms through the landscape. Disturbance events can have severe consequences for these ecosystems and the entire downstream freshwater network and require more attention in the context of global change-induced increases in weather extremes. Here we show that extreme rain events (floods) cause cyclic successions in microbial communities and the planktonic food web of a small forest pond. We analyzed the dynamics of nutrients and the entire plankton community during two flood events and subsequent quasi-stable conditions. Floods induced a repeated washout of resident organisms and hundred-fold increases in nutrient load. However, within two weeks, the microbial community recovered to a pre-disturbance state through four well-defined succession phases. Reassembly of phyto- and especially zooplankton took considerably longer and displayed both repetitive and adaptive patterns. Release of dissolved nutrients from the pond was associated with inflow rates and state of community recovery, and it returned to pre-disturbance levels earlier than microbial composition. Our study exemplifies extraordinary compositional and functional resilience of small waterbodies and presents the detailed mechanism of the underlying processes.

---

Nutrients, materials, and organisms transferred from terrestrial environments to freshwater networks play a crucial role in global biogeochemical cycles^1–3^. The shoreline of small lakes and ponds represents the largest surface area among all inland waters and thus serves as an important terrestrial-aquatic interface^3–5^. Uptake, transfer and release of nutrients occurring in the food web of small waterbodies thus have considerable consequences for the entire downstream aquatic network^6–9^. At the same time, small water volume and the tight linkage to the surrounding land render these waterbodies highly vulnerable to fluctuating hydrological conditions. Extreme rain events (ERE) shorten water retention times (WRT) to values characteristic for streams, cause washout of planktonic organisms, and bring extreme loads of nutrients and organic matter from flooded watersheds^9–13^. Such strong disturbances on planktonic communities can result in delaying, arresting or diverting seasonal succession from its usual pattern^14, 15^.

Intensification in strength and frequency of EREs due to an ongoing and future climate change is predicted for many areas worldwide, including Central Europe^16–18^. Although terrestrial-aquatic boundaries have a crucial influence on the entire landscape as highly responsive hotspots of organic matter transformation, energy and nutrient flow^19^, flood-induced effects on nutrient budgets, composition and function of plankton communities as well as on food web structure remain largely unknown. While communities of small water bodies cannot possess resistance to EREs due to direct washout effects, their resilience capacity^20^ is of high interest, raising an intriguing question to which extent these ecosystems are able to absorb EREs, or if such disturbances force them beyond their resilience limits towards new alternative steady states^21, 22^. To increase the predictability of consequences caused by EREs, extensive high temporal resolution studies along with an application of integrative and dynamic concepts extendable from population to meta-ecosystem levels are necessary.

In our study, we examined the planktonic community organization from viruses to zooplankton and fluxes of major elements in a small humic waterbody with strong aquatic-terrestrial coupling. Our high-resolution *in situ* sampling campaign during two extreme rain events and subsequent quasi-stable conditions addressed two fundamental questions: (I) What are the patterns in responses of community compositions and functions to ERE disturbances? (II) What is the role of small headwater ecosystems in the transfer of nutrients and organisms across the terrestrial-freshwater interface at stable and extreme hydrological conditions?

We could show that although EREs caused a washout of resident organisms, the microbial community recovered within two weeks in four well-defined succession phases. While reassembly of phyto- and especially zooplankton took considerably longer, dissolved nutrients, which were enhanced hundred-fold during the floods, returned to pre-disturbance levels even earlier than microbes. We thus conclude that the microbial community and the aquatic food web, in general, are not resistant but extraordinary resilient and show a fast recovery after disturbances by EREs. Moreover, the microbial community of small ponds plays a crucial role as a sink of soluble nutrients that would otherwise be exported to the downstream network. This important function was rapidly re-established after the EREs, emphasizing high functional resilience of such ecosystems.

## Results and discussion

### General overview of sampling site and hydrological conditions

For our study, we have chosen Jiřická Pond^23, 24^ (Extended Data Fig. 1) located in the headwaters of Malše River (CZ). It is a small humic water body strongly influenced by forested landscape representing a freshwater ecosystem typical for Central and Northern Europe (Extended Data Text 1). The pond was studied during highly-dynamic hydrological period controlled by precipitation (May 5^th^ to June 27^th^, 2014). To unveil ERE-associated changes in the plankton community and to estimate terrestrial influences on organismic dynamics, two sites (pond epilimnion and stream – a main tributary of the pond) were sampled three times per week. Sampling started one month after the snow-melt at low inflow rates (0.04 to 0.27 m^3^ s^−1^) resulting in a WRT in the pond surface layer between 6.7 and 11.1 days. The precipitation rates did not exceed 11.3 mm d^−1^ before May 16^th^, when continuous rainfall for three days introduced 79.5 mm of water and caused the first flood event in the pond catchment (ERE1). Another heavy rainfall yielded precipitation of 112.3 mm and resulted in the second ERE (ERE2) equivalent to a 5-years flood around May 28^th^. Both EREs severely decreased the WRT of the pond surface layer to 0.46 d (ERE1) and 0.44 d (ERE2) and increased flow rates of the stream to 2.2 m^3^ s^−1^ (ERE1) and 2.5 m^3^ s^−1^ (ERE2) (Fig. 1a-b). That strongly affected physicochemical parameters at both sampling sites (Extended Data Text 1), e.g. seasonally rising water temperatures and stable pH values were interrupted by two pronounced ERE-related drops (Fig. 1c-d).

**Figure 1:**
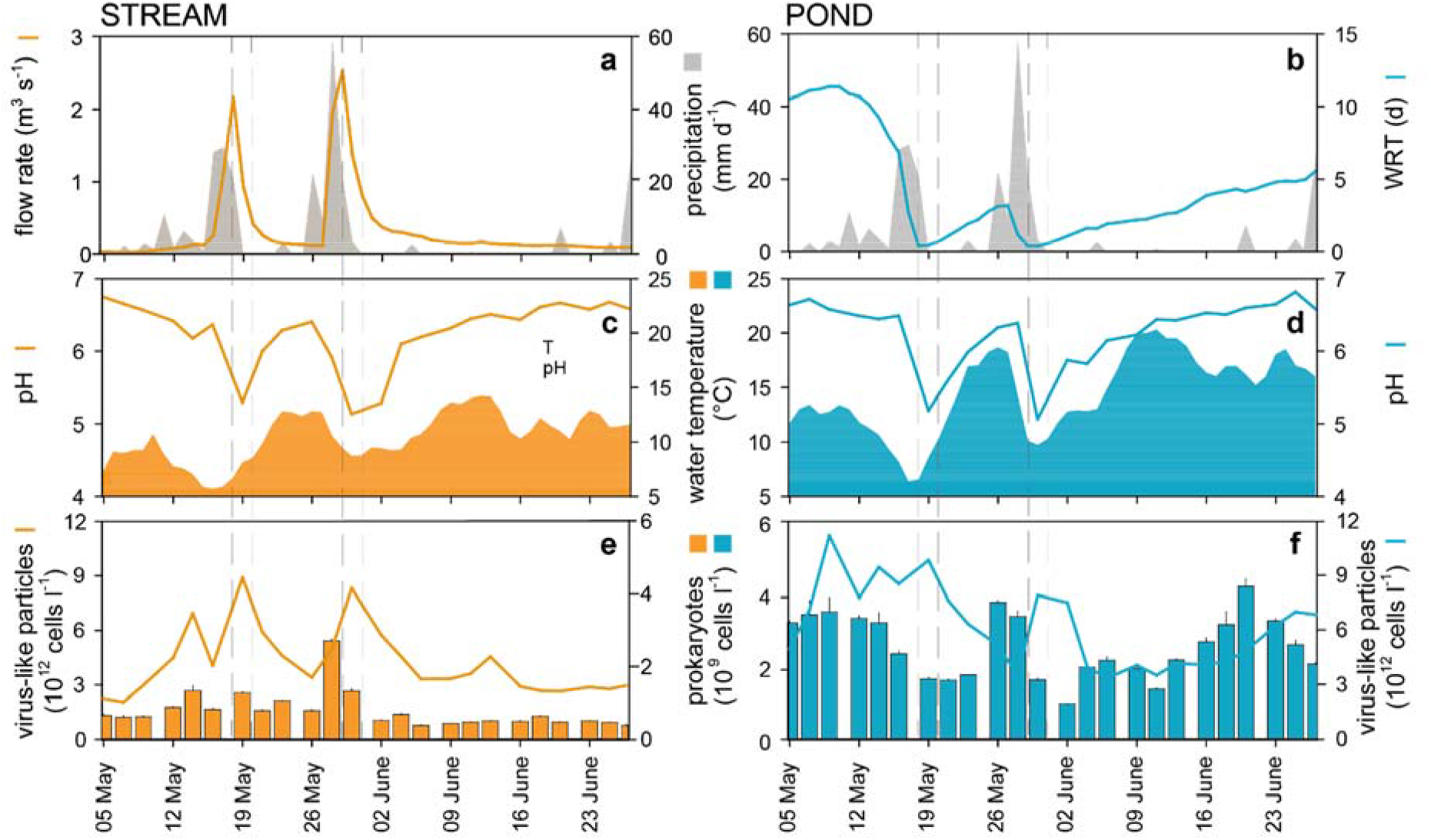
Hydrological, physicochemical and microbial parameters measured in the stream (left: **a, c, e**) and pond (right: **b, d, f**). **a, b**: Precipitation (grey shade) in the watershed of Jiřícká pond and stream flow rate (**a**) as well as water retention time (WRT) at 0.5m depth in the pond (**b**). **c**, **d**: Water temperature and pH. **e**, **f**: Abundances of prokaryotic cells (counted in duplicates, bars represent ranges) and virus-like particles (VLP). The dashed lines indicate two EREs observed during the sampling campaign.

### Flood-caused microbial dynamics in the pond

Dynamics of bacteria, which represent numerically the most resistant part of the pond plankton to washout-effects, was studied with two approaches: (i) 16S rRNA gene amplicon sequencing to access bacterial community diversity and (ii) catalyzed reporter deposition *in situ* fluorescence hybridization (CARD-FISH) for accurate enumeration of the 20 most abundant bacterial groups. Distance matrices calculated for data obtained with these methods displayed high agreement (r^2^=0.63, p<0.0001, two-sided Mantel test) and revealed that bacterial community composition underwent pronounced shifts after both EREs, but rapidly cycled back to the pre-disturbed stage during subsequent developments (Fig. 2).

**Figure 2:**
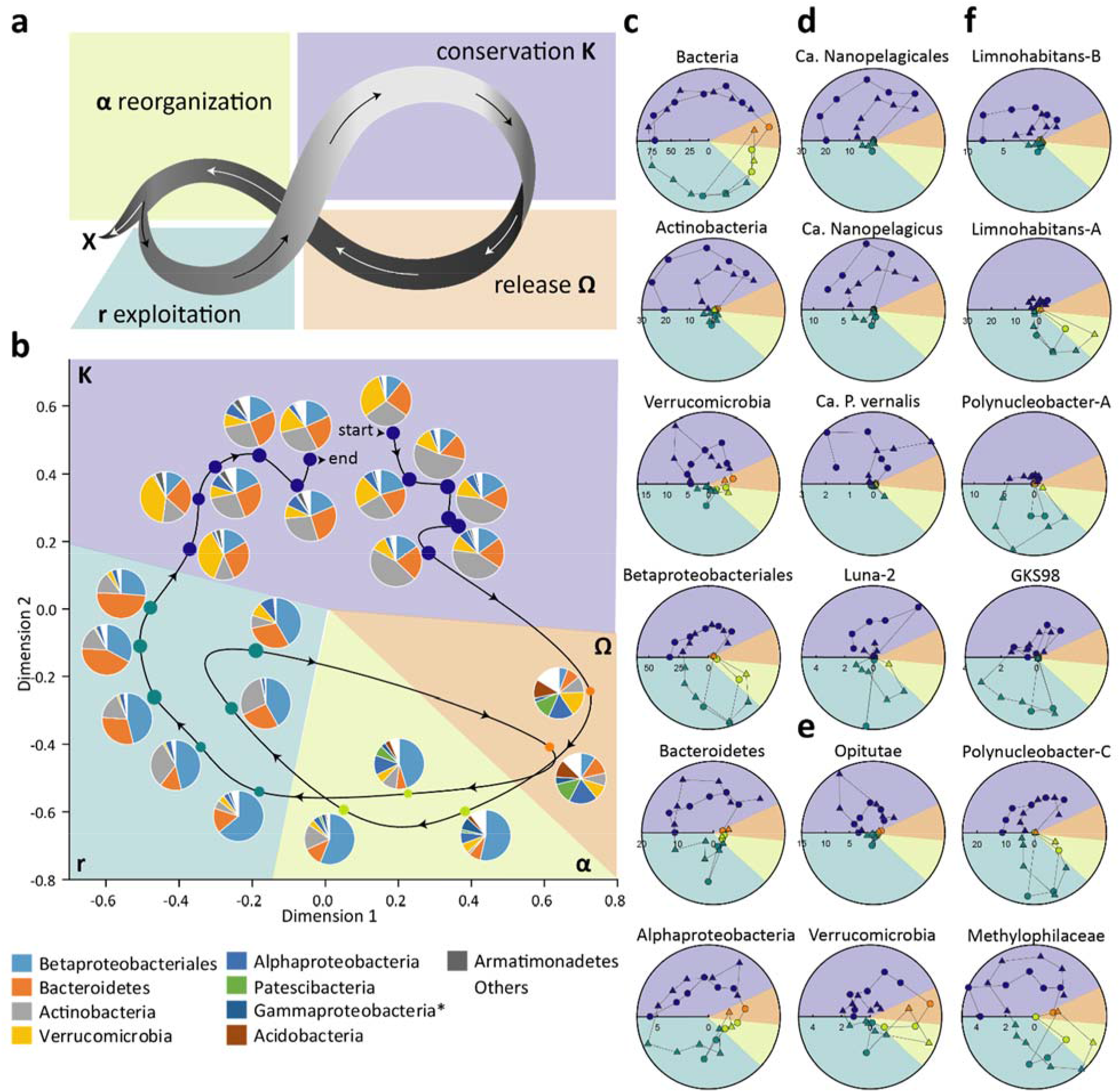
Bacterial succession in the pond following the adaptive cycle. **a**: Representation of the adaptive cycle^22^; α and Ω phases comprising the back-loop, r and K phases represent the fore-loop of the cycle. X indicates the path to an alternative development **b**: Multidimensional scaling (MDS) based on 16S rRNA gene amplicons from pond bacterial communities (Kruskal's stress: 0.17; Anosim test for 4 phases: R=0.95, P=0.0001). A line connects samples following chronological order. The diameters of the circles are proportional to the Pielou’s evenness of the samples (varying between 0.3 and 0.4). Pie charts depict the bacterial diversity at the class-phylum level. Letters and associated colors visualize the phases of observed succession. *: Gammaproteobacteria are represented without Betaproteobacteriales, which are shown separately. ‘**K**’ and violet color indicate conservation-phase, ‘**Ω**’ and orange – collapse and release phase, ‘**α**’ and light-green – reorganization-phase, ‘**r**’ and aquamarine – exploitation-phase. **c-g**: Radial plots depicting percentages of bacterial groups obtained by CARD-FISH (in % of hybridized bacteria). Radial axes represent percentages. Angular axes represent time. Colors of segments and symbols correspond to succession-phases: The time series start with circles: from violet (first K-phase) to aquamarine (first r-phase). Subsequent samples in triangles represent the second Ω-phase (orange), second α-phase (light-green), second r-phase (aquamarine), and final K-phase (violet).This graph design enables easy comparisons between samples taken in the same phase. **c**: Relative abundances of general bacterial groups. **d**: Relative abundances of selected Actinobacteria groups. **e**: Relative abundances of Verrucomicrobia groups. **f**: Relative abundances of selected Betaproteobacteriales groups.

According to agglomerative clustering analysis (Extended Data Fig. 2a), four distinct phases could be distinguished in this succession pattern, which in their sequence and characteristics followed the course of an “adaptive cycle” (Fig. 2a) that was repeated twice during our entire high-frequency sampling (Fig. 2b). This “adaptive cycle”, a concept initially developed to describe community organization and dynamics in complex ecosystems^22^, consists of four well-defined phases (Fig 2a). Two of those, namely so-called ‘r’ (growth and exploitation) and ‘K’ (conservation) phases, were adopted from succession ecology^25^ and represent the slow fore-loop of the cycle characterized by biomass accumulation and an increase in connectedness within the ecosystem. The phases ‘Ω’ (collapse and release) and ‘α’ (reorganization) are parts of the fast back-loop of the cycle that results either in a reorganization of the system and escape towards an alternative steady state or in the repetition of the previous cycle. These twice-repeated phases were also consistent with trends in several chemical parameters and food web organization and are described in details as follows:

**Conservation phase (K)** was observed at the beginning and the end of the sampling campaign, i.e. the first six and last six samplings (Fig. 2b-e, Extended Data Fig. 3). During these periods, the pond was characterized by a WRT longer than 3.9 days and high consumption rates of soluble inorganic nutrients (Fig. 3). Up to 60% of the dissolved reactive phosphorus (DRP) and up to 100% of nitrate and ammonia transported by the inlet were consumed during that period in the pond indicating that plankton growth was severely nitrogen limited (Extended Data Fig. 2). As expected, a relatively low bacterial diversity but high community evenness (Extended Data Fig. 3) were characteristic for these resource-limited conditions^22^. The majority of amplicon sequence variants (ASVs) detected during K phase (65.5%) had closest relatives reported from aquatic environments and consisted of a few dominant taxa typical for humic lakes^26^ (Actinobacteria, Verrucomicrobia, Bacteroidetes and Betaproteobacteriales; Fig. 2c). Highest proportions of K phase specific ASVs (Extended Data Fig. 4a) were affiliated with ‘Ca. Nanopelagicales’ (Actinobacteria) and the family Puniceicoccaceae (Verrucomicrobia, Opitutae), which have small-sized cells and genomes^27, 28^, and can be considered as typical *K*-strategists. According to the *r*/*K*-selection ecological concept, those are slow-growing organisms, for which a stable environment selects due to limited resources^29, 30^. During the K phase, the organization of the microbial food web, phytoplankton and higher trophic levels in the pond was most complex (Fig. 4, Extended Data Fig. 6), corresponding to the expected high connectedness in the system^22^ (Fig. 2a). This resulted in the highest bacterial loss rate due to increased protistan bacterivory indicating a severe top-down control of the bacterial community (Extended Data Fig. 7). That provided conditions favorable for grazing-resistant microbes, e.g. ‘*Ca*. Nanopelagicales’, protected from protistan grazers due to their small cell size^27, 31^. Thus, both top-down control and *K*-selection contributed to “species sorting” by habitat conditions^32^, which dominated in microbial community assembly during the K phase.

**Figure 3:**
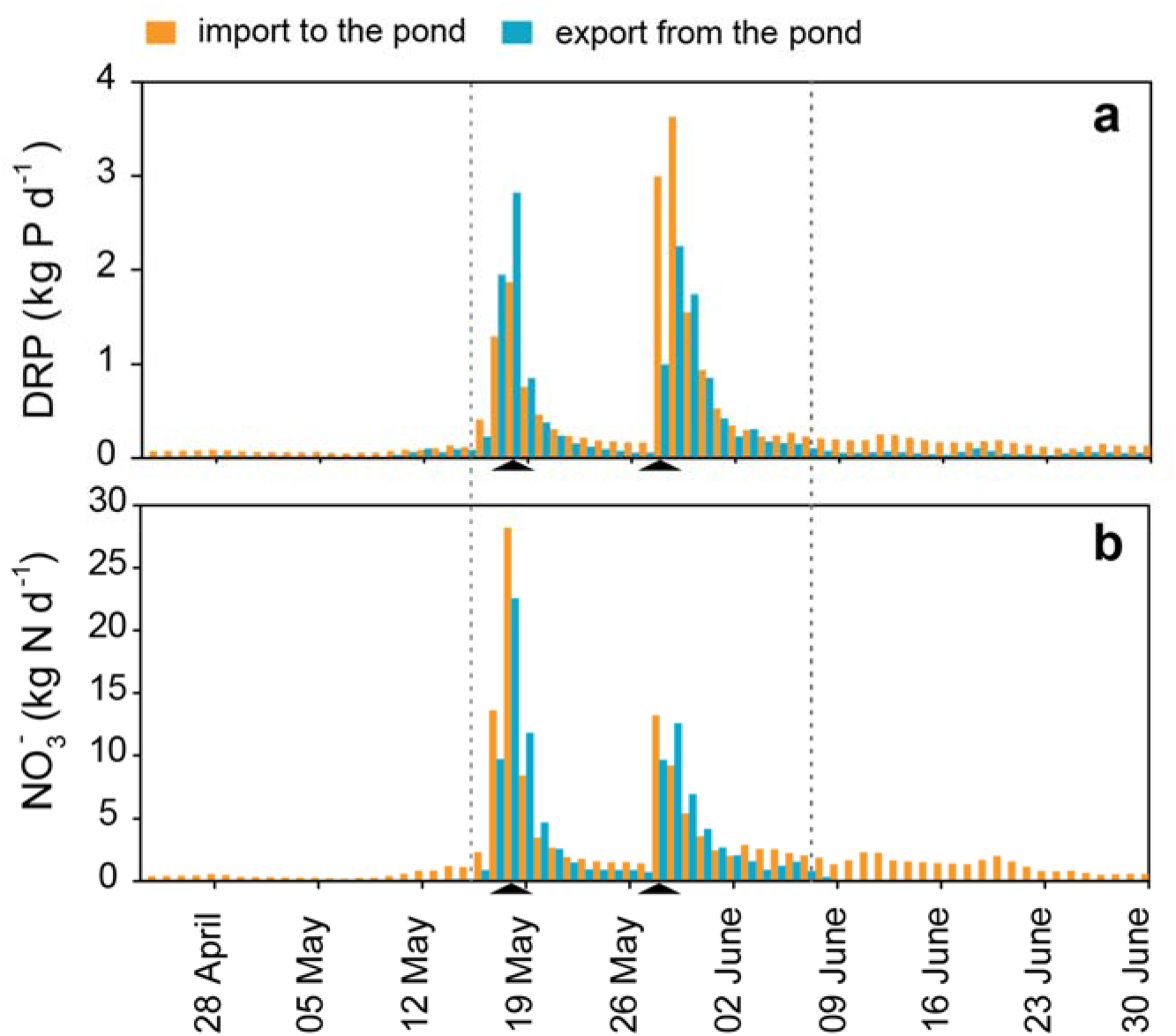
Daily import and export rates of **a:** dissolved reactive phosphorus (DRP), **b**: nitrate. Import to the pond is depicted in orange, export from the pond is depicted in blue. Dotted lines are separating three equal periods (23 days each) used for the calculation of exports and imports at different hydrological conditions. Black triangles indicate EREs.

**Figure 4:**
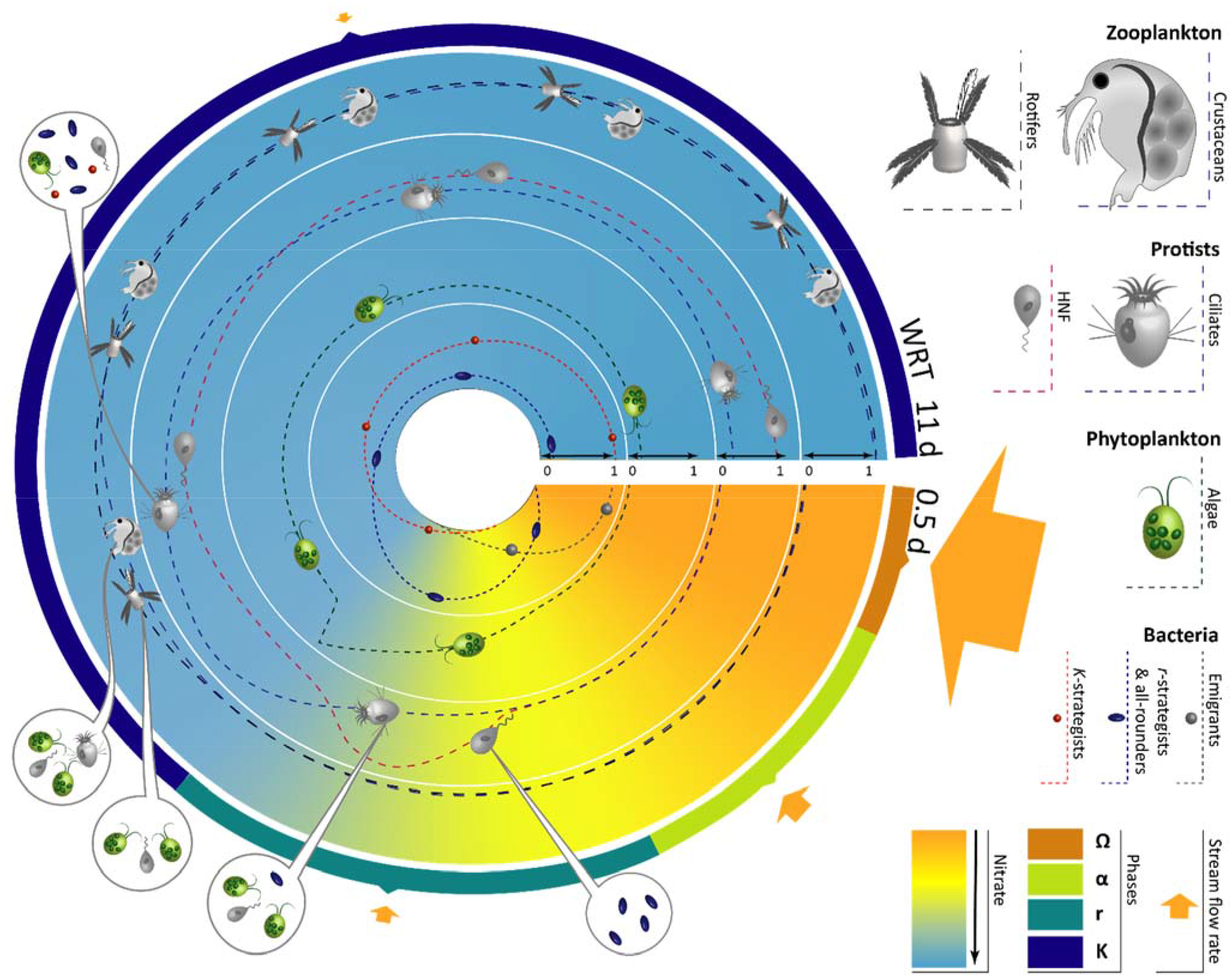
Food web structure of the Jiřická Pond plankton community during flood-caused cyclic succession. Different phases are indicated in an external circle with colors: violet – ‘K’ conservation phase, orange – ‘Ω’ collapse and release phase, light green – ‘α’ reorganization phase, and aquamarine – ‘r’ exploitation phase. Relative abundances (where ‘1’ corresponds to maximum abundances) of different plankton groups are depicted in the four inner circles (from outside to inside): Zooplankton is represented by rotifers (dominated by *Polyarthra sp*.) and crustaceans (dominated by *Eubosmina* sp.). Protists are represented by heterotrophic nanoflagellates (HNF) and ciliates. Food preferences of zooplankton and protists at different phases are indicated by bubbles. Phytoplankton is represented by summarized dynamics of all algal species. Bacterioplankton is divided into three categories: emigrants, *r*-strategists & all-rounders, and *K-*strategists. Nutrient gradient based on nitrate concentrations (0.00 to 0.14 mg N l^−1^) is displayed by different background colors, stream flow rates are indicated by different sized arrows.

**Mass effects-driven collapse and release phase (Ω)** consists of two samples collected during the EREs (Extended Data Fig. 2a). They were characterized by short WRT (<0.46 d) and up to two-fold increased concentrations of DOC and dissolved nutrients (Fig. 3, Extended Data Fig. 8). The phase displayed low prokaryotic densities but yielded the highest diversity and lowest evenness of the bacterial community (Extended Data Fig. 4), resembling flood-associated microbial dynamics in subsurface ponds^33, 34^. The proportion of ASVs reported from aquatic habitats decreased to less than half (18%), with only 3% being associated with lake bacteria. A high proportion of soil-related ASVs (46%) in the pond indicated a tight terrestrial-aquatic coupling and an important role of “mass effects”, a term used in metacommunity theory for emigration processes^32^. Emigrant phylotypes (from the terrestrial surrounding) within Verrucomicrobia, Alphaproteobacteria, Acidobacteria, Solirubrobacteraceae, Omnitrophicaeota and Patescibacteria (Extended Data Fig. 5c) displayed short-lived peaks, but did not persist in the epilimnion during the subsequent stabilizing hydrological conditions, revealing high “colonization resistance”^35^ of the pond (Extended Data Fig. 5c).

Virus-like particles (VLPs) in both pond and stream had local maxima during EREs (Fig. 1e-f), which was most likely connected to the introduction of soil VLPs and/or resuspension of VLPs from the stream sediment. In contrast, larger plankton groups (protists, phytoplankton and zooplankton) were almost entirely washed-out from the pond (Extended Data Fig. 5a-c).

**Reorganization phase (α)** comprises three samples collected between the third and fifth day after the ERE peaks. This phase was characterized by a still low WRT in the pond (<2.0 d) and enhanced DOC and nutrients concentrations (Fig. 1, Extended Data Fig. 8). Instead of soil-associated emigrant ASVs detected in Ω-phase, we observed high read numbers of ASVs associated with the stream (Extended Data Fig. 5d), e.g. representatives of *Duganella*, *Rhodoferax*, *Flavobacterium* and *Pseudomonas* lineages. At the same time, we detected several pond-specific ASVs, e.g. representatives of *Polynucleobacter* cluster C^36^ (PnecC), ‘*Ca*. Methylosemipumilus turicensis’^37^, as well as a pond specific ASV of *Limnohabitans* lineage LimA (Extended Data Fig. 5e-f). The observed occurrence patterns exemplify an evident reorganization in assembly mechanisms of the pond bacterial community, namely a decrease of mass effects and the appearance of species sorting selecting during this phase for so-called *r-*strategists, fast-growing organisms able to rapidly exploit resources and to colonize unstable environments^29, 30, 38^. Epilimnetic populations of protists, phyto- and zooplankton still remained at negligible levels.

**Exploitation-phase (r)** contains seven samples (two after ERE1 and five after ERE2). In the course of this stage, chemical and hydrological characteristics, as well as consumption rates of soluble inorganic nutrients changed to levels comparable to stable conditions due to increasing exploitation. Linkages between bacterial communities of pond and stream were reduced (Extended Data Fig. 5c-e); as with increasing WRT, stream relevant ASVs were replaced by pond specific ones. Betaproteobacteriales displayed maxima in CARD-FISH and sequence read proportions accounting for up to 57.5% of all bacteria (Fig. 2c). Among them, several ASVs were mainly restricted to the exploitation phase: the majority of *Limnohabitans* lineage LimA, previously reported as EREs associated copiotrophs^39^, seemed to represent typical *r*-strategists; while an ASV of PnecA (*Polynucleobacter rarus)* could be favored by a decrease in pH^40^. On the other hand, ASVs of PnecC and ‘*Ca*. Methylosemipumilus turicensis’ showed persistent read and cell count proportions also in the α and K phases (Fig. 2f, Extended Data Fig. 5f). Both taxa are small to medium-sized bacteria with passive lifestyles based on the utilization of photodegradation products or C1 compounds^37, 41^. This seemingly successful strategy renders them to abundant all-rounders in the described ecosystem. During the r phase, also fast-growing algae and heterotrophic nanoflagellates (HNF) attained their maxima, while ciliates, rotifers and crustaceans displayed their peaks later^23^ (Fig. 4, Extended Data Fig. 6). Uncontrolled growth of HNF together with nutrient limitation resulted in the collapse of bacterial *r-*strategists, which thereafter were replaced by grazing resistant *K*-strategists, e.g. ‘*Ca*. Nanopelagicales’ and Bacteroidetes^27, 38, 42^. This transit from r to K phase completed the observed succession cycle.

### The ERE-induced succession of the entire planktonic community

The bacterial community in the pond revealed an unexpected resilience in response to EREs and returned to the initial steady state through the above-described well-determined pathway within 16 days after ERE2. The fifty most common pond ASVs displayed highly repetitive appearance patterns in the two loops and were mainly restricted to one or two distinct phases. Though, we missed the recovery of three ASVs well represented during the first conservation phase (one representative of Verrucomicrobiae and two of Bacteroidetes, Extended Data Fig. 5a), not even a single emigrant ASV could establish successfully in the pond after EREs. The succession of pond eukaryotes generally followed patterns outlined in the PEG (Plankton Ecology Group) model^15, 43^ tightly related to the organisms’ size and growth strategy. Fast-growing bacterial *r-*strategists were followed by bacterivorous HNF, having comparable growth potential^44^ as these prey bacteria. In parallel, we observed a rapid increase of small green flagellates (*Chlamydomonas* spp., Extended Data Fig. 6). With a short delay of 7 days, the growth of these edible algae, as well as HNF, became top-down controlled by the subsequently recovered ciliate community (Fig. 4, Extended Data Fig. 6–7), represented mainly by predatory prostome ciliates^45^. After the decline of edible HNF and algal species, more omnivorous ciliates were detected in the pond. This led to an increase in ciliate bacterivory during the second conservation phase (Extended Data Fig. 7) and higher loss rates of Verrucomicrobia and Bacteroidetes (Extended Data Fig. 6d, g), which appear to be less susceptible to HNF grazing, but highly vulnerable to omnivorous ciliates (e.g. *Halteria* sp. *Rimostrombidium* spp.). Similar to bacteria, eukaryote dynamics were driven by resource-limitations and disturbances but also by top-down control via food web structure, dominated at different successional stages by HNF, ciliates, crustaceans or copepods (Fig. 4, Extended Data Fig. 5, 9). As expected for dynamic waterbodies, zooplankton in the pond was greatly dominated by rotifers (average ratio to crustaceans 126:1^23, 46^ Extended Data Fig. 6, 10). After EREs, reassembled populations of cladocerans and rotifers displayed a higher diversity, but did not reach the pre-disturbance densities because of the establishment of a predatory copepod population (*Mesocyclops leuckarti*^23^, Extended Data Text 1). Recovery of this species in early summer represents a well-known seasonal development^47^. Temporal patterns were also observed in the phytoplankton dynamics, e.g. during the second conservation phase, *Chlamydomonas* spp. persisted in the phytoplankton with Cryptophytes and Chrysophytes, typical representatives of spring assemblages^31^ (Extended Data Fig. 6, 9). Seasonal dynamics of phyto- and zooplankton seemed to be accompanied by stochastic processes in the post EREs reassembly (back-loop of the adaptive cycle) that resulted in alternations in their composition (Extended Data Fig. 9–10). Surprisingly, the reorganization of higher trophic levels from ciliates to cladocerans showed a limited influence on bacterial community composition observed in both K phases.

### Segregation between stream and pond

Most physicochemical characteristics in both stream and pond displayed comparable dynamics tightly linked to EREs (Extended Data Fig. 3 and 8). Significant differences between the stream and pond were observed only for temperature, which was higher in the pond (Mann-Whitney rank test: P<0.001) and soluble fractions of nitrogen and phosphorus, which were higher in the stream (Mann-Whitney rank test: P<0.001 for dissolved phosphorus (DP), and nitrate; P=0.022 for ammonia; two-tailed *t*-test: *t*(46)=-0.8, P=3‧10^−9^ for DRP). Despite similar chemistry, constantly low WRT did not allow the establishment of a stable plankton community in the stream. In result, we could not identify typical stream eukaryotic plankton populations due to their low densities, or recognize clear succession patterns caused by EREs even in bacterial populations (Extended Data Fig. 12). High mass effects resulted in 26% of soil associated bacterial ASVs in the stream even during a low hydrological regime. Despite high stochasticity in the community assembly, high nutrient contents and low grazing pressure provided favorable conditions for *r*-strategists in the stream. Such ASVs were detectable in the stream at all non-EREs conditions, but in the pond exclusively during the reorganization period that increased the similarity between both sides (Extended Data Fig. 2c, 5e, 11). However, a WRT of two days was sufficient to separate inlet and pond microbial communities and to prevent an overlap of the most abundant phylotypes (Extended Data Fig. 2, 12). This notion demonstrates the compositional discontinuity between the hydrologically connected stream and pond ecosystems, and limited influence of mass effects on pond community assembly during stable hydrological conditions, i.e. the K phase.

### Transfer of nutrients through the pond

Low initial flow rates (0.07±0.03 m^3^ s^−1^; AV±SD) at the beginning of the sampling campaign (April 23^th^ – May 16^th^, 2014) were associated with very low loads of dissolved nutrients from the watershed (Extended Data Fig. 3). The average daily export of nitrate and DRP from the pond amounted to 0.05±0.07 kg and 0.04±0.03 kg, respectively. Altogether, during this period, 1.05 kg of nitrate N (8% of the import) and 0.96 kg DRP (48% of the import) were exported from the system. Loads of soluble nutrients increased dramatically with rain events. Daily export of nitrate and DRP reached up to 22.6 kg and 2.6 kg, respectively. In 23 days, covering both EREs and community recovery periods, 103.7 kg of nitrate N and 14.6 kg of DRP were released to the downstream network. The subsequent equivalent period was characterized by slightly elevated flow rates and loads from the watershed (0.13±0.03 m^3^ s^−1^; 31.8 kg nitrate N; 4.1 kg DRP), but also by an already recovered microbial community and hence low nutrient export (0.7 kg nitrate N; 1.5 kg DRP). This indicates that ponds substantially control the nutrient release from the watershed, as nitrate export remained in the same range during pre and post EREs’ periods, while nitrogen consumption in the pond differed more than three times following EREs (Extended Data Fig. 3).

### Conclusions

In this study, we present the detailed mechanism underlying functional and compositional resilience of the small pond to EREs and connect the flow of nutrients, bacterioplankton dynamics and reorganization of higher trophic levels to the entire ecosystem development. The backbone of community recovery is represented by the twice-repeated adaptive cycle in bacterioplankton composition. Four well-defined phases of this cycle reflected the transition from the dominance of mass effects during and shortly after the EREs to the prevalence of species sorting in community assembly mechanisms during conservation phases. Within this successional development, we report remarkable fast return of nutrient export rates to pre-disturbance levels in just nine days, rapid reestablishment of the pro- and eukaryotic microbial community within 16 days, and slightly longer reassembly of phyto- and zooplankton, which was to a greater degree influenced by the seasonal succession of different taxa. Thus, our high-resolution sampling campaign revealed that small waterbodies have the potential of surprisingly fast and almost complete compositional and functional recoveries after ERE disturbances. However, as the frequency of extreme rain events will increase in the future, it remains questionable to which extent this resilience can be challenged and how possible alternations in the composition could affect ecosystem function and provided services. Knowledge on functional and compositional resilience of small waterbodies, representing the most abundant freshwater habitats of Earth, is of high importance for sustainable management and for predicting the impact of future climate change on their role in transformation and control of organic matter and nutrient flow through the landscape.

## Material and methods

### Study Site

Jiřická Pond (Pohořský Pond) is located in Novohradské Mountains at an elevation of 892 m a.s.l. (48.6189N, 14.6731E) and was formed by damming of the Pohořský stream in the 18^th^ century. Jiřická is a small (0.04 km^2^), shallow (4 m max. depth) humic pond, the catchment (12.5 km^2^) is comprised to a high extend by forest but also by pastures and raised peat bogs ^24^. The intensive sampling campaign of the pond and the main tributary of Jiřická (stream) took place between May 5^th^ and June 27^th^, 2014, with samplings three times per week (Mon, Wed and Fri; total number of samplings: 24). Samples for chemical and biological analyses were taken with a Friedinger sampler at 0.5 m depth at the deepest point of the pond. The stream was sampled 500 m upstream from the pond.

### Meteorological and hydrological characteristics

Meteorological parameters, including the level of precipitation, were monitored with the meteorological station M4016 (Fiedler-Mágr, Czech Republic) installed near the pond during the study period. The water level of the inlet was measured three times per week, and the flow rates were calculated based on the relation between water level and discharge calibrated at different flow regimes using handheld Acoustic Doppler Velocimeter (FlowTracker, Son Tek, USA). Water age at 0.5 m was calculated using the CE-QUAL-W2 model of reservoir hydrodynamics^48^.

### Enumeration of planktonic organisms

Samples for microbial analyses (2 l) were split into subsamples that were either immediately fixed on-site or transported to the laboratory within two hours. For enumeration of virus-like particles (VLPs), one ml of water samples from both sites was fixed with glutaraldehyde (1% final concentration) for 10 min, shock-frozen in liquid nitrogen, and transported in liquid nitrogen to the lab, where samples were stored at −80°C. Enumerations of VLPs were done using an inFlux V-GS cell sorter (Becton Dickinson) as previously described^49^. For bacterial enumeration, water subsamples were fixed with formaldehyde on site (2% final concentration). In the lab, duplicates of 1-2 ml were stained with DAPI and filtered onto black 0.2 μm pore-size polycarbonate membranes. The bacterial abundance was determined via epifluorescence microscopy^50^. Numbers of heterotrophic nanoflagellates (HNF) and ciliates were estimated in a similar manner from Lugol-formaldehyde-thiosulfate fixed subsamples^51^. For this purpose, 5-30 ml were stained with DAPI and filtered onto black 1.0 μm pore-size membrane. Protozoan bacterivory was estimated using fluorescently labelled bacteria (FLB)^52^ prepared from a mixture of cultures of small rods of *Limnohabitans* lineage LimC^44, 53^ and two undescribed strains of *Polynucleobacter* lineage PnecC. The sizes of FLB reassembled the typical size class distribution of the bacterioplankton in the pond. HNF and ciliate abundances and FLB uptake rates were determined in short-term FLB direct-uptake experiments, with ciliates being determined to genus level^54^. To estimate total protozoan grazing, we multiplied the average uptake rates of HNF and ciliates with their *in situ* abundance. Phytoplankton samples were preserved with acid Lugol solution and stored in a dark box at room temperature. Species were enumerated with the Utermöhl method^55^ using the microscope Olympus IMT2. The mean algal cell dimensions for biovolume calculation were obtained using the approximation of cell morphology to regular geometric shapes^56^.

Zooplankton was sampled once a week. Crustaceans were sampled by vertical hauls using an Apstein plankton net (200 μm mesh) from the entire water column^57^ (4 m). Rotifers were sampled from the whole water column using a plastic tube of the appropriate length. A total of 55 l of water was subsequently filtered using a 35 μm net. The zooplankton samples were preserved in 4% formaldehyde, and species abundances were determined microscopically.

### Nucleic acids sampling, isolation and processing

Microbial biomass was collected on Sterivex filter units (polyethersulfon membranes, 0.22 μm pore size, Millipore, USA) by filtering approximately 600 ml with a sterile syringe (n=48). Additional samples for RNA isolation were prepared in the same manner every Wednesday (n=14). The units were sealed, flush-frozen in liquid nitrogen and transported in liquid nitrogen to the lab, and stored at −80°C until further processing. Nucleic acids (DNA and RNA) were isolated using phenol-chloroform-isoamyl alcohol extraction method according to a previously described protocol^58^. After washing with 70% ethanol, nucleic acids were briefly dried, dissolved in 20 μl of nuclease-free water and stored at −80°C. For cDNA synthesis, DNA was degraded in the DNA/RNA extract by twice repeated incubation with Turbo DNase at 37°C for 45 min, followed by addition of DNase inactivation reagent according to the Turbo DNA-free Kit protocol (Ambion, USA). Thereafter, 20 to 50 ng of RNA were mixed with 1.5 μl primer mix (Random Primers 6, New England BioLabs, MA, USA) in RNA free water (final volume 10.5 μl) and incubated for 5 min at 70°C. Reverse transcription was performed with the Array Script enzyme (Ambion, USA) according to the manufacturer’s instructions at 42°C for 2 h and terminated by enzyme deactivation at 95°C for 5 min.

### Amplicon sequencing and analysis

Primers F27 and R519 were used to amplify the V1-V3 region of the 16S rRNA gene. A single-step PCR using HotStarTaq Plus Master Mix Kit (Qiagen Inc, Valencia, CA, USA) was conducted as follows: 94°C for 3 minutes, followed by 28 cycles of 94°C for 30 seconds; 53°C for 40 seconds and 72°C for 1 minute; after which a final elongation step at 72°C for 5 minutes was performed. After PCR, all amplicon products were mixed in equal concentrations and purified using Ampure beads (Agencourt Bioscience Corporation, MA, USA). The amplicons were sequenced using the Roche 454 GS FLX Titanium platform at MR DNA laboratory (Shallowater, TX, USA) with an average sequencing depth of 50000 raw reads per sample.

Demultiplexed reads were processed using DADA2 package version 1.12 in R^59, 60^. The primer free sequences were trimmed to 350 bp length according to the quality report. The values recommended for 454 data were used for the denoising, namely homopolymer gap penalty ‘-1’ and increased to 32 band size. Chimeric amplicon sequence variants (ASVs) were detected and removed with de novo algorithm. The remaining unique ASVs were taxonomically assigned using naive Bayesian classifier method and trained Silva SSU database release 132^61, 62^. Variants affiliated to mitochondria and plastids were excluded. Two samples with read numbers <10 000 were removed from analysis (stream DNA from 25.06.2014 and stream cDNA from 11.06.2014). The final number of reads in samples ranged from 19390 to 87185; the dataset was rarefied to 19390 reads per sample prior to diversity and statistical analyses. From the pond samples, the 50 most common ASVs were selected according to the percentages of reads per sample for a graphical representation of read abundance dynamics. Since these phylotypes were not detected in the EREs samples, the same procedure was done additionaly for most frequent ASVs in the EREs samples (>0.3% of read number in the sample; n=14). The next relatives of these sequences and of 50 most common ASVs from stream samples were assigned with BLAST 2.2.21 against Silva SSU database release 132^61, 63^ and a bootstrapped Randomized Axelerated Maximum Likelihood (RAxML) tree was constructed (GTR-GAMMA model, 100 bootstraps^64^). The isolation source information for the next relatives with identity scores >97% was collected from NCBI using reutil package^65^ in R^60^ and manually classified to water vs. soil related environments and others.

### Clone libraries of 23S rRNA genes and probe design for CARD-FISH

Clone libraries of 23S rRNA genes were constructed for three dates: 12.05.2014; 26.05.2014 and 02.06.2014. DNA extracts and primers 113f and 2744r_mod^39^ (Extended Data Table 1) were used for the amplification of 23S rRNA genes using a Taq polymerase (Top-Bio, Czech Republic). After initial denaturation step (3 minutes at 94°C), 28 cycles were performed at the following conditions: 30 seconds at 94°C, 30 seconds at 50°C and 3 minutes at 72°C; that was followed by the final elongation step at 72°C for 10 minutes. The purified PCR products were cloned into *Escherichia coli* using PCR Cloning^plus^ kit, (QIAGEN, Germany). Plasmid inserts of colourless colonies were amplified by PCR with M13F and M13R primers^66^, PCR products were purified and sequenced with M13F primer. Trimmed sequences were taxonomically classified with the ARB software and SILVA LSU database release 132^61, 67^. PCR products affiliated with bacterial taxa of interest (i.e., Betaproteobacteriales and Actinobacteria, n=68), were additionally sequenced with four primers (Extended Data Table 1), partial sequences were assembled using Geneious R9 software (www.geneious.com), aligned with the SINA web-aligner^68^ and imported to ARB for tree reconstruction and CARD-FISH probe design. As the general primer 1358f^39^ was not matching all Actinobacteria (Extended Data Table 1), an additional primer targeting Actinobacteria (1367f) was designed in ARB using the tools probe_design and probe_check. Alignments of 68 almost full-length sequences (>2000 nt) and their closest relatives were manually improved and two RAxML trees (GTR-GAMMA model, 100 bootstraps^64^ were constructed for *Limnohabitans* spp. (Extended Data Fig. 13) and Actinobacteria affiliated with ‘*Ca*. Nanopelagicales’ and the luna2 lineage (Extended Data Fig. 14), respectively. Probe design for the LimB cluster of *Limnohabitans* spp. and the luna2 cluster of Actinobacteria was done with the tools probe_design and probe_check. Three competitor probes were designed for probe LimB-23S-920 and two competitor probes and three helper oligonucleotides were designed for probe luna2-23S-1239 to further increase mismatch discrimination and rRNA accessibility^69^. Additionally to 23S rRNA targeted probes, we also designed and modified a set of probes targeting the 16S rRNA genes of Verrucomicrobia as an online check with SILVA^62^ revealed that the general probe ver47^70^ targets only a part of freshwater Opitutales (79%, Extended Data Fig. 15). We modified this probe in a way that it discriminated against this order (probe ver46: 0.1% coverage of Opitutales), and designed an additional probe that targets 97.1% of all available sequences of Opitutales (probe opitu-346; Extended Data Fig. 15). All probes were checked *in silico* for specificity and coverage using the TestProbe function of SILVA^62^. Probes were further tested *in silico*^71^ and in the laboratory with different formamide concentrations in the hybridization buffer until stringent conditions were achieved (Extended Data Table 2).

#### Catalyzed Reporter Deposition Fluorescent in situ Hybridization (CARD-FISH) analysis

For CARD-FISH analyses, formaldehyde-fixed samples from the pond epilimnion (n=24) were filtered within 10 hours after fixation onto white polycarbonate membrane with 0.2 μm pore size (47 mm diameter, Millipore, USA). Filters were stored at −20°C prior to subsequent analysis. The filter sections were used for specific hybridization of 24 bacterial groups with horseradish peroxidase labelled oligonucleotide-probes (Extended Data Table 2), and subsequent amplification with fluorescein labelled tyramides as previously described^72^. Proportions of CARD-FISH positive signals were evaluated with an epifluorescence microscope (Olympus BX-53F) for at least 500 DAPI-stained cells per probe.

### Physicochemical characteristics

Water temperature was monitored during the entire sampling period at the pond and stream sampling sites at hourly intervals with temperature data loggers (Tidbit, Onset, USA) and averaged for daily values. Samples for other physicochemical analyses were collected from pond and stream sites at every sampling in prewashed and rinsed polyethylene terephthalate sampling bottles and were delivered to the laboratory in thermos-boxes within two hours. Chlorophyll *a* concentration was determined spectrophotometrically after extraction with acetone^73^. Acid neutralizing capacity (determined by Gran titration) and pH were measured by Radiometer TIM 865 station (Hach, Germany). Conductivity was evaluated using inoLab®Cond 720 (WTW, Germany). Aliquots of samples were filtered through 0.45 μm nylon membrane filters (Fisher Scientific, USA) for ion chromatography and silica determination. Dissolved reactive silica was determined by the molybdate method^74^. Concentrations of cations (Na^+^, K^+^, Ca^2+^, Mg^2+^, NH_4_^+^) and anions (Cl^−^, SO_4_^2–^, F^−^, NO ^−^, NO ^−^) were determined by ion chromatography (Dionex IC25, USA). All following chemical analyses, except for total phosphorus, were performed after filtration of water samples through 0.4 μm glass fiber filter (GF-5, Macherey-Nagel, Germany). Dissolved reactive phosphorus (DRP) was determined by the molybdate method^75^, and total (TP) and dissolved phosphorus (DP) with a modification of this method with sample pre-concentration and perchloric acid digestion^76^ yielding detection limits of 1 and 0.5 μg l^−1^, respectively. DOC was analyzed as a non-purgeable organic carbon by catalytic combustion at 680°C (Elementar, Germany), then the sample was acidified with HCl (1 M) to pH below 4 and air purged for three min prior to the analysis. The detection limit was ~0.05 mg l^−1^. DN concentrations were obtained using a vario TOC cube (Elementar, Germany), nitrogen was determined as NO by infrared-(NDIR) detector below the ppb level.

Molecular weight distribution of organic compounds was determined for all samples collected after May 16^th^ by high-performance size exclusion chromatography (Dionex DX320, USA) with a photodiode array detector (ICS-3000 PDA) with detection at 210, 225, and 254 nm. A PL-aquagel-OH 20 column (7.5 mm × 30 cm and a particle size of 5 μm; Agilent, USA) was used for size exclusion. For each run, 200 μl of the sample was injected onto the column with phosphate buffer (0.1 M NaCl, pH 6.8) as an eluent, and the buffer flow rate was set at 0.5 ml min^−1^. The column void volume and total permeation volume of the column were determined using Blue Dextran and acetone, respectively. Polyethylene Glycol/Oxide standards (Agilent, USA) were used as molecular mass calibration standards (molecular weights: 420, 615, 1480, 3870, 7830, 12300, and 21300 Da). Low molecular weight organic (carboxylic) acids (lactic, acetic, formic, butyric, pyruvic, adipic, benzoic, tartaric, maleic, ascorbic, fumaric, citric, and iso-citric acids) were determined by capillary ion chromatography (ICS-5000, ThermoFisher, USA) equipped with IonPac AS11-HC-4 μm and IonPac AG11-HC-4 μm columns with conductivity detection. A standard capillary EGC KOH with multistep nonlinear gradient was used for elution with concentrations from 0.5 mM to 110 mM. The range of calibration for acetic and formic acid was 50 – 1000 μg l^−1^ while it was 10 – 200 μg l^−1^ for other acids.

### Data analyses

Diversity estimators and indexes for rarefied data of 16S rRNA gene amplicons and Anosim statistics with 9999 permutations were calculated with the vegan package^77^ in R. Clustering analysis, multidimensional scaling, calcultion of Bray-Curtis dissimilarity matrixes, two-tailed Mantel tests (Pearson) 95% confidence interval were performed using XLSTAT 14 (Addinsoft). *t*-tests or alternatively Mann-Whitney rank tests (for not normal distributed data e.g. TP, DP, amonia, nitrate nitrite, pH, temperature) on 95% confidence interval were performed with SigmaPlot 12.5 after normality Shapiro-Wilk tests.

The nutrient balances represent the differences between values obtained from a transfer model (theoretical values calculated for the nutrient transport to epilimnion based on nutrient concentrations measured in the stream and hydrological parameters) and values measured in the pond epilimnion. The transfer of different compounds from the inlet to the epilimnion of Jiřická Pond was modeled using the hydrodynamic and water quality model CE-QUAL-W2^48^. For modelling purposes, the water body was separated into 12 segments, and all calculations were done for 0.5 m depth layers^48^. The import and export of nutrients were calculated by multiplying the average daily flow data by the nutrient concentrations.

## Data availability

16S rRNA and rDNA amplicon data from this study were deposited to NCBI under BioProject PRJNA547706, BioSamples SAMN11974970-11975031. Sequences of 23S rRNA genes obtained from clone libraries were submitted to NCBI under accession numbers MN565597-MN565664.

## Acknowledgements

The authors would like to thank all scientific and technical staff supporting the team during the exhausting but also exiting sampling campaign, especially Radka Malá and Hana Kratochvílová. The study was supported by projects from the Czech Science Foundation: project 20-23718Y awarded to TS, project 13-00243S awarded to KŠ, project 19-00113S awarded to PP, projects 19-23469S and 20-12496X awarded to MMS. HPG was supported by the German Ministry of Education and Science (BMBF) in the frame of the BIBS project (TP 7: 01LC1501G).

## Author contributions

TS, MMS and KŠ designed the study, collected and enumerated bacterial and protist samples. TS contributed to CARD-FISH analysis, performed bioinformatics analysis of amplicon sequences and statistical analyses of data, interpreted the data, designed the graphs and wrote the manuscript. MMS designed the CARD-FISH probes, contributed to CARD-FISH analysis, prepared the phylogenetic trees and participated in data interpretation and manuscript writing. PP collected and analyzed chemistry samples, contributed to interpretation of data and manuscript revision. PZ and JN collected and analyzed samples for phytoplankton and contributed to interpretation of data and manuscript revision. HPG was responsible for the isolation of the nucleic acids and preparation of the amplicon libraries and participated in data interpretation and manuscript writing. JS collected and analyzed zooplankton data, contributed to interpretation of data and manuscript revision. JH collected and analyzed meteorological and hydrological data, calculated the transfer models and nutrient balances, and contributed to interpretation of data and manuscript revision. KŠ analyzed protist community-related parameters and contributed to interpretation of data and manuscript writing.

## Additional information

**Extended Data Fig. 1:**
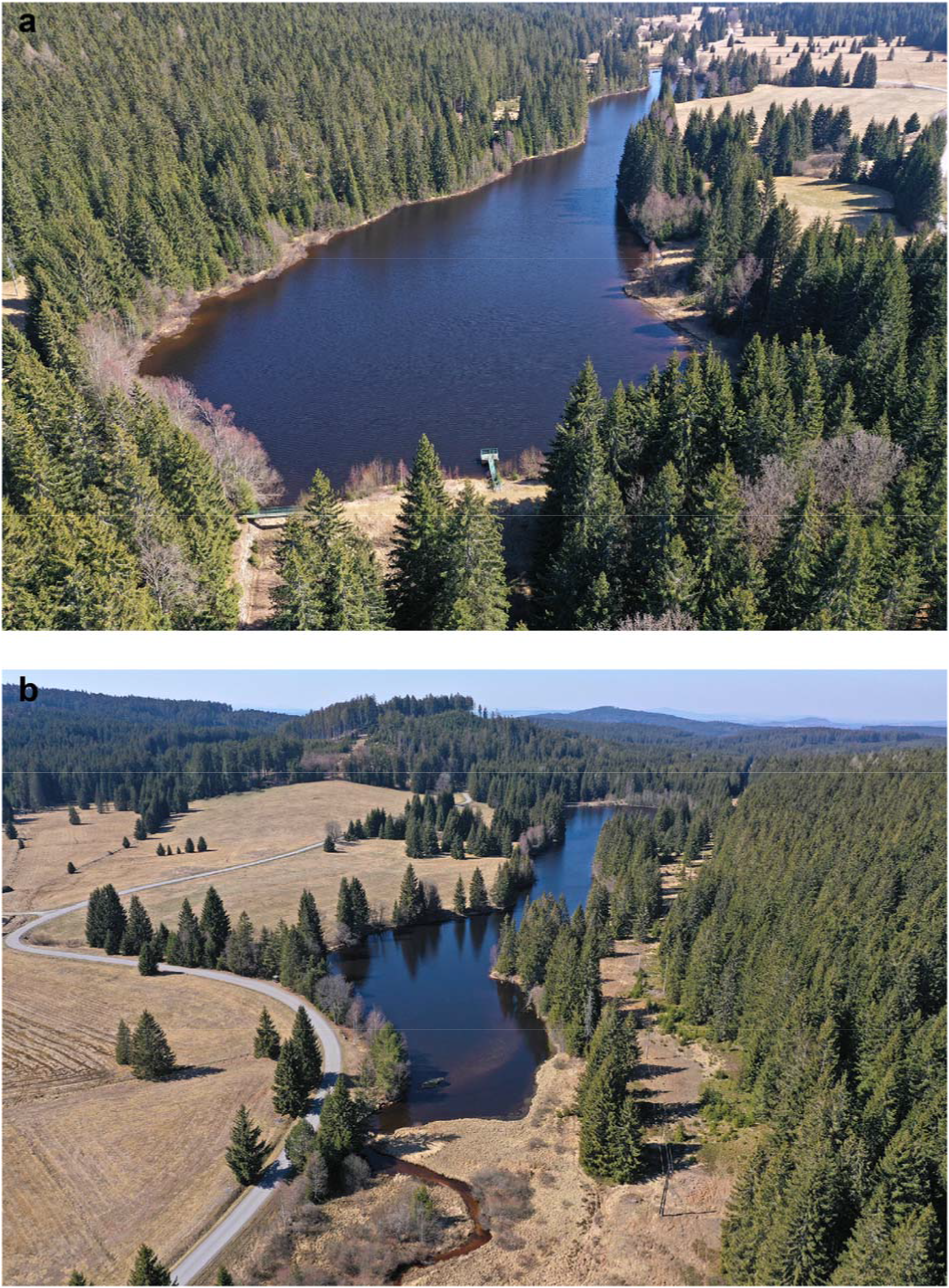
Aerophotographs of Jiřická Pond. **a:** Dam site; **b:** Stream site. Photographs were kindly provided by Petr Znachor.

**Extended Data Fig. 2:**
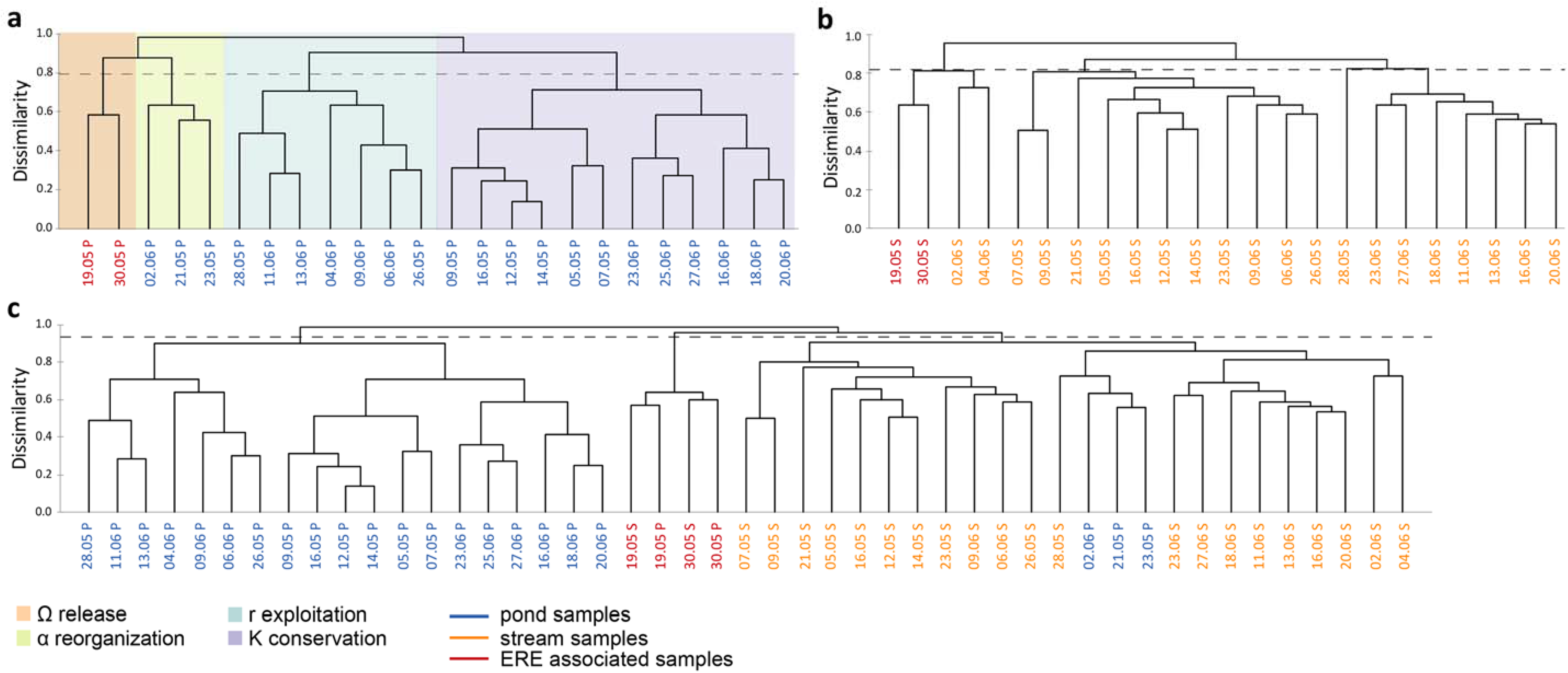
Agglomerative hierarchical clustering (AHC) based on Bray-Curtis dissimilarity matrix of 16S rDNA amplicon data. Dashed lines show the level of truncation. Sample names are composed of the date and the first letter of the sampling site (P – pond, S – stream). Samples from the pond are depicted in blue, samples from the stream in orange, samples collected during EREs are shown in red. **a:** AHC of pond samples. Colored background areas highlight clusters, which defined four phases of the succession cycle: violet – ‘K’ conservation phase, orange – ‘Ω’ release phase, light green – ‘α’ reorganization phase, and aquamarine – ‘r’ exploitation phase. **b:** AHC of stream samples. **c:** AHC of all stream and pond samples.

**Extended Data Fig. 3:**
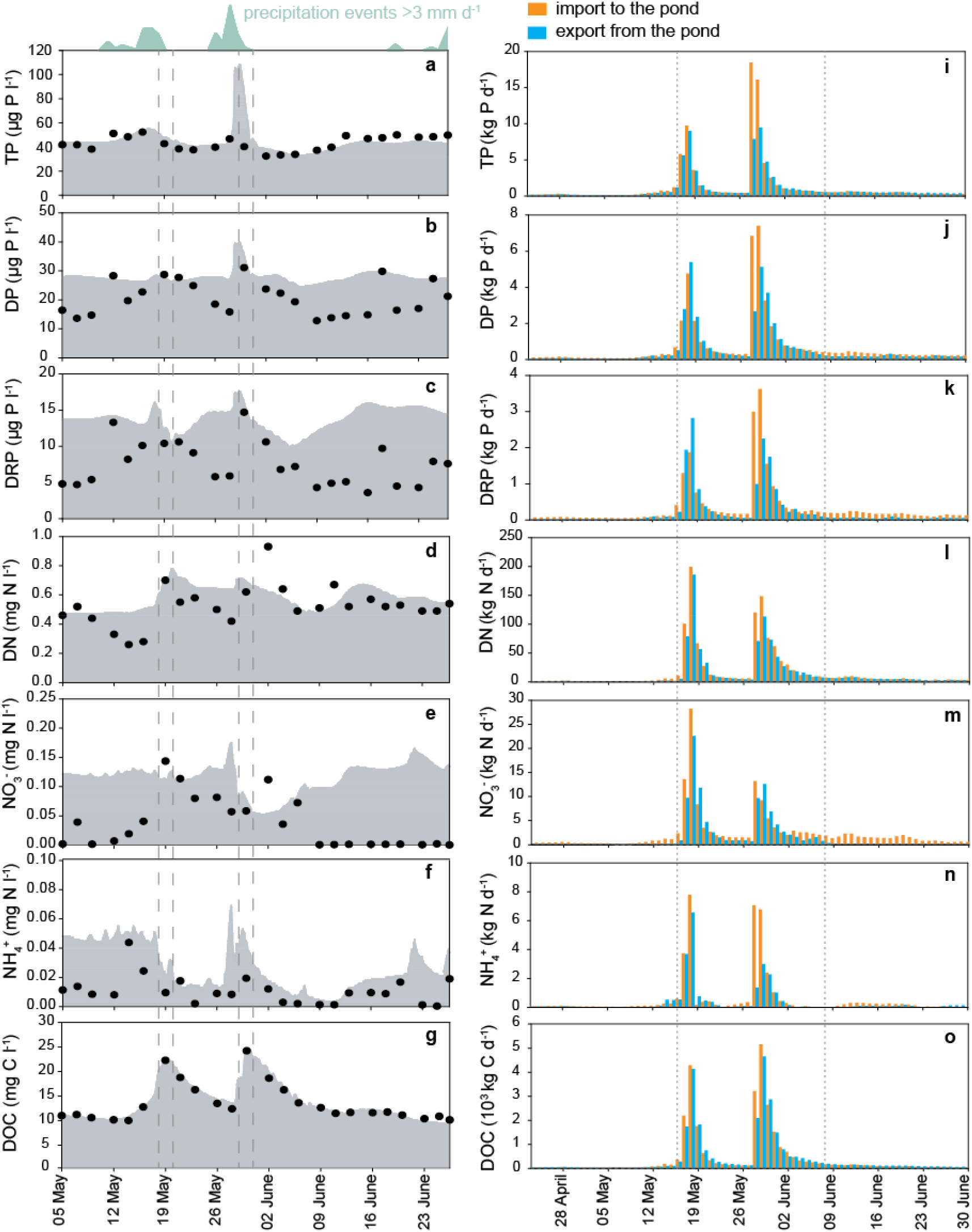
Mass balances of different chemical compounds in Jiřická Pond. **a-g**: Black circles represent values of compounds measured in the pond. Grey surfaces represent theoretical hydrological transport of compounds from the inlet (calculated with CE-QUAL-W2 Hydrodynamic and Water Quality Model using settings for conservative tracer). Dashed lines indicate two flood events observed during the sampling campaign. Precipitation events >3 mm d^−1^ are depicted in aquamarine above panel **a**. **a-c:** Phosphorus fractions. TP, total phosphorus; DP, dissolved phosphorus; DRP, dissolved reactive phosphorus. **d-f:** Nitrogen fractions. DN, dissolved nitrogen. **g**: Dissolved organic carbon (DOC). **i-o:** Daily import and export rates of different compounds. Dotted lines separate three equal periods of 23 days each. Import to the pond is shown in orange, export from the pond in blue. **i-k:** Phosphorus fractions. TP, total phosphorus; DP, dissolved phosphorus; DRP, dissolved reactive phosphorus. **l-n:** Nitrogen fractions. DN, dissolved nitrogen. **o**: Dissolved organic carbon (DOC).

**Extended Data Fig. 4:**
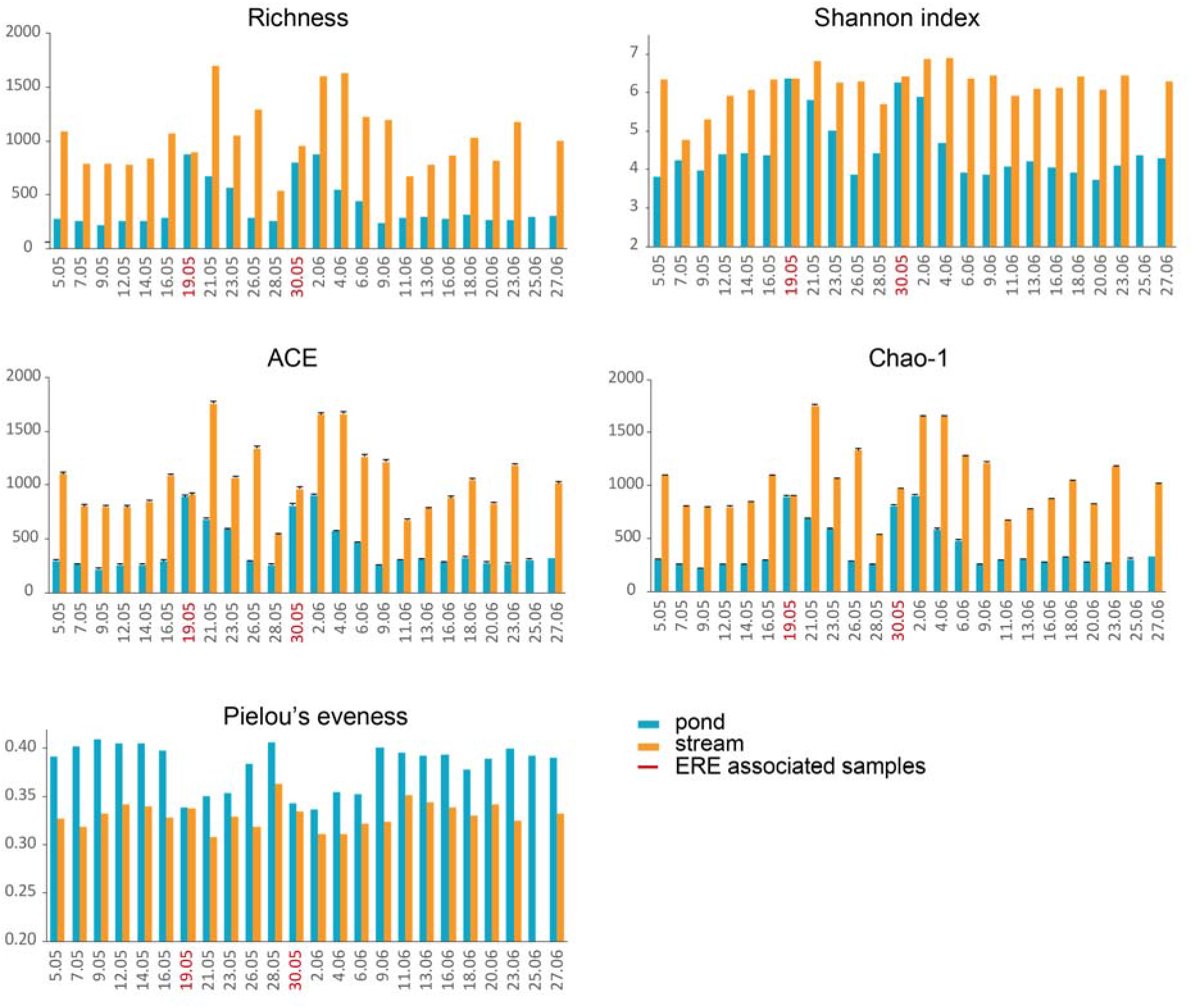
Community diversity and evenness parameters observed at pond and stream sites during the sampling campaign. Error bars represent SEs. Red font indicates dates of EREs.

**Extended Data Fig. 5:**
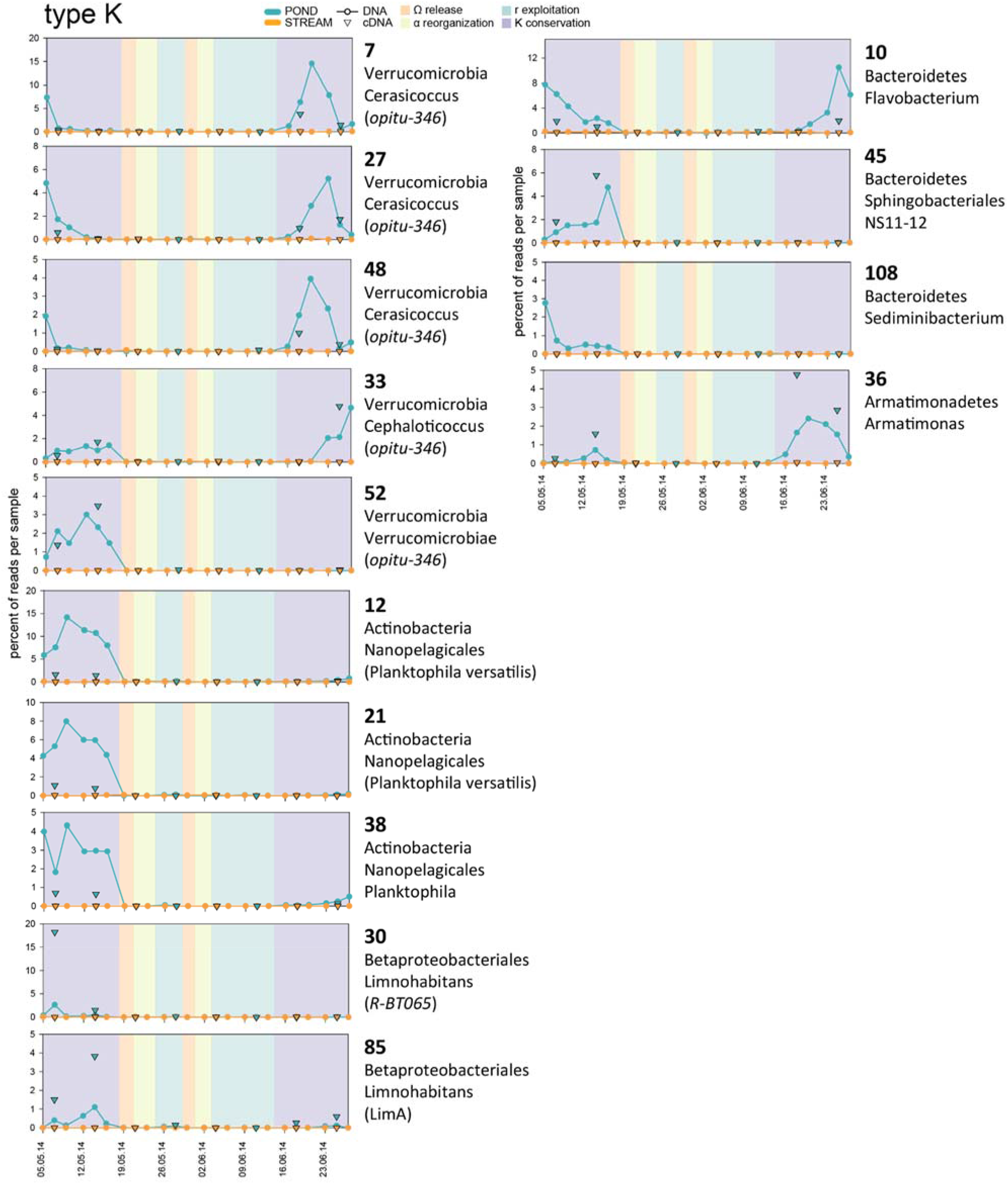
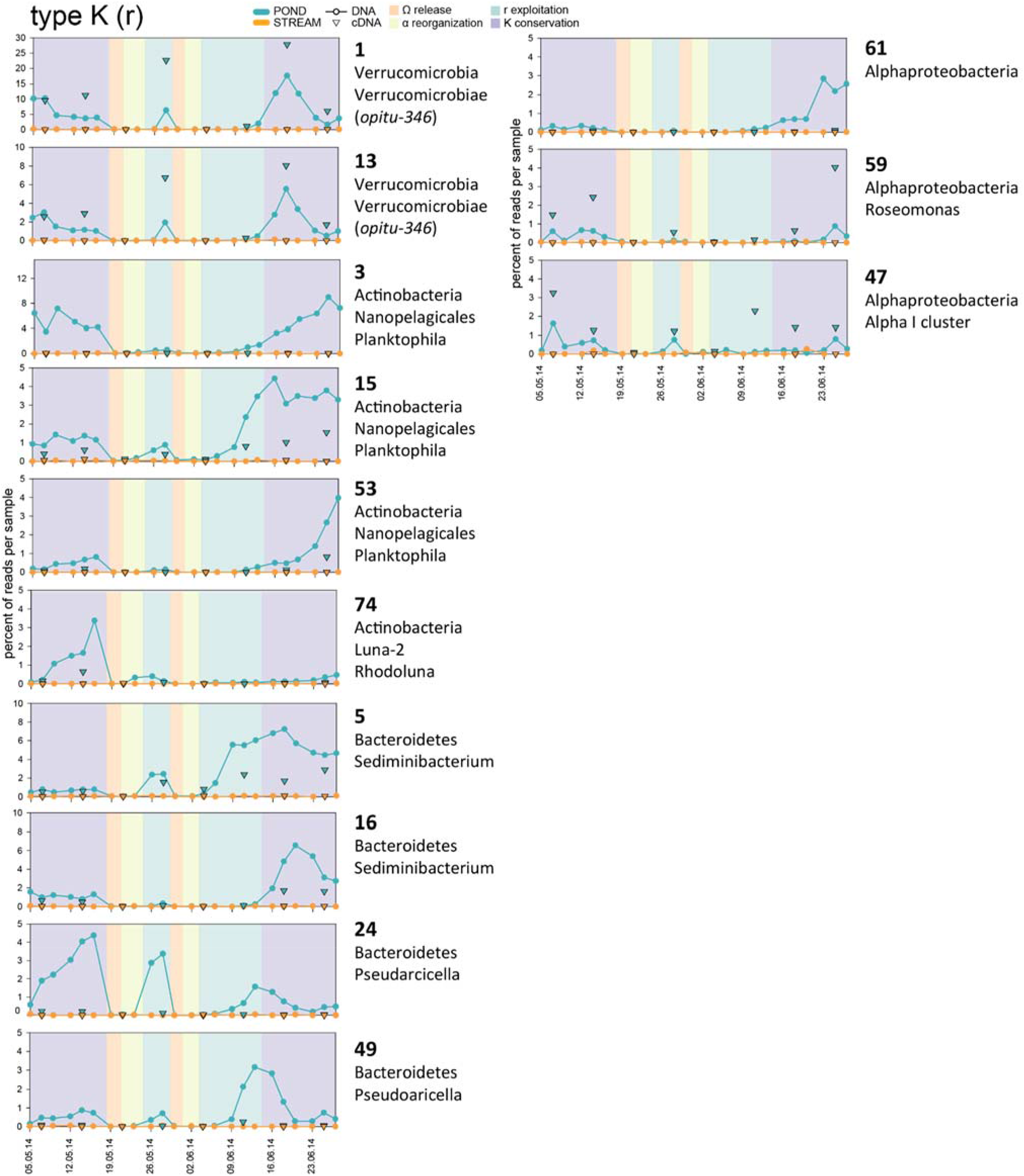
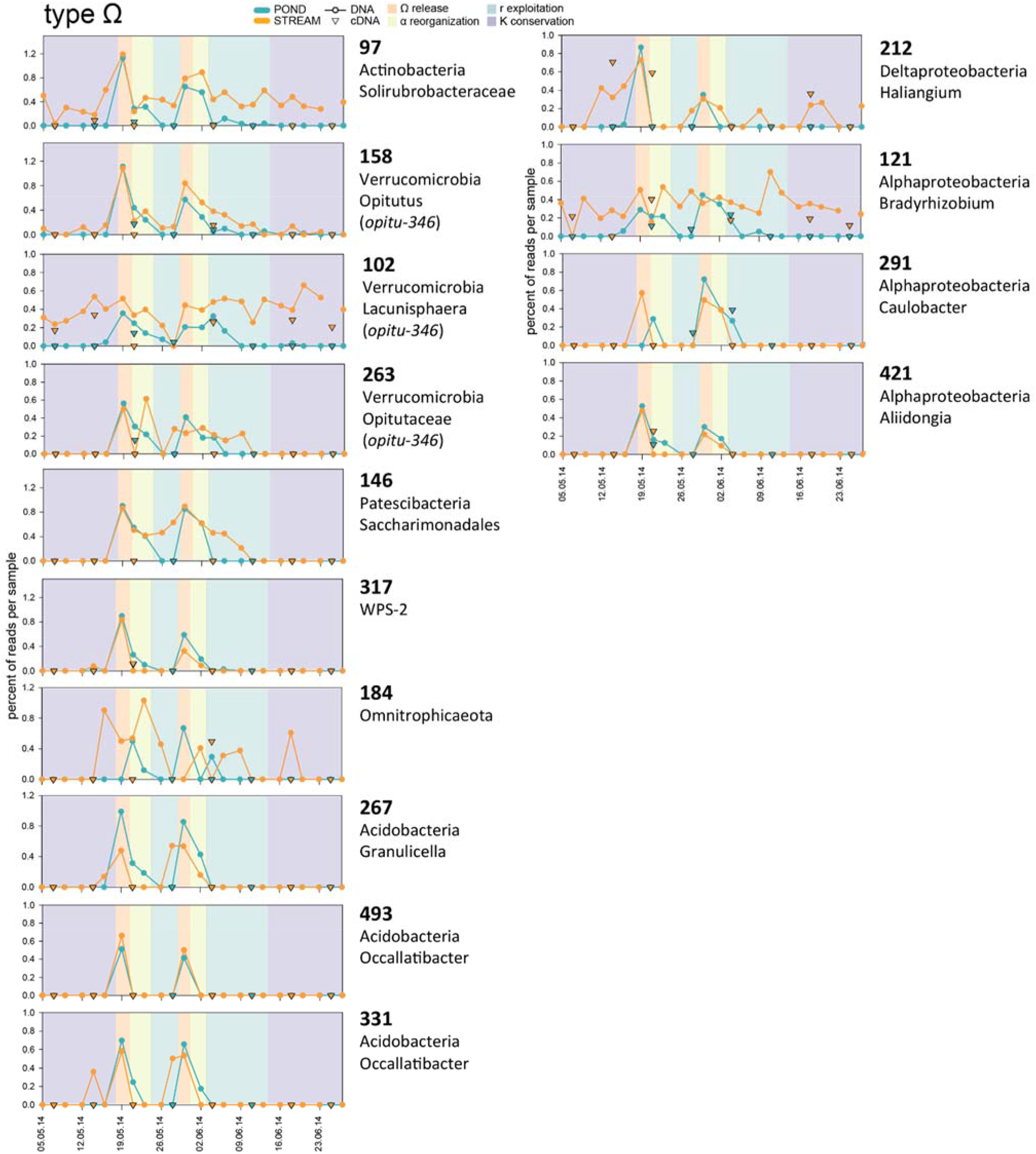
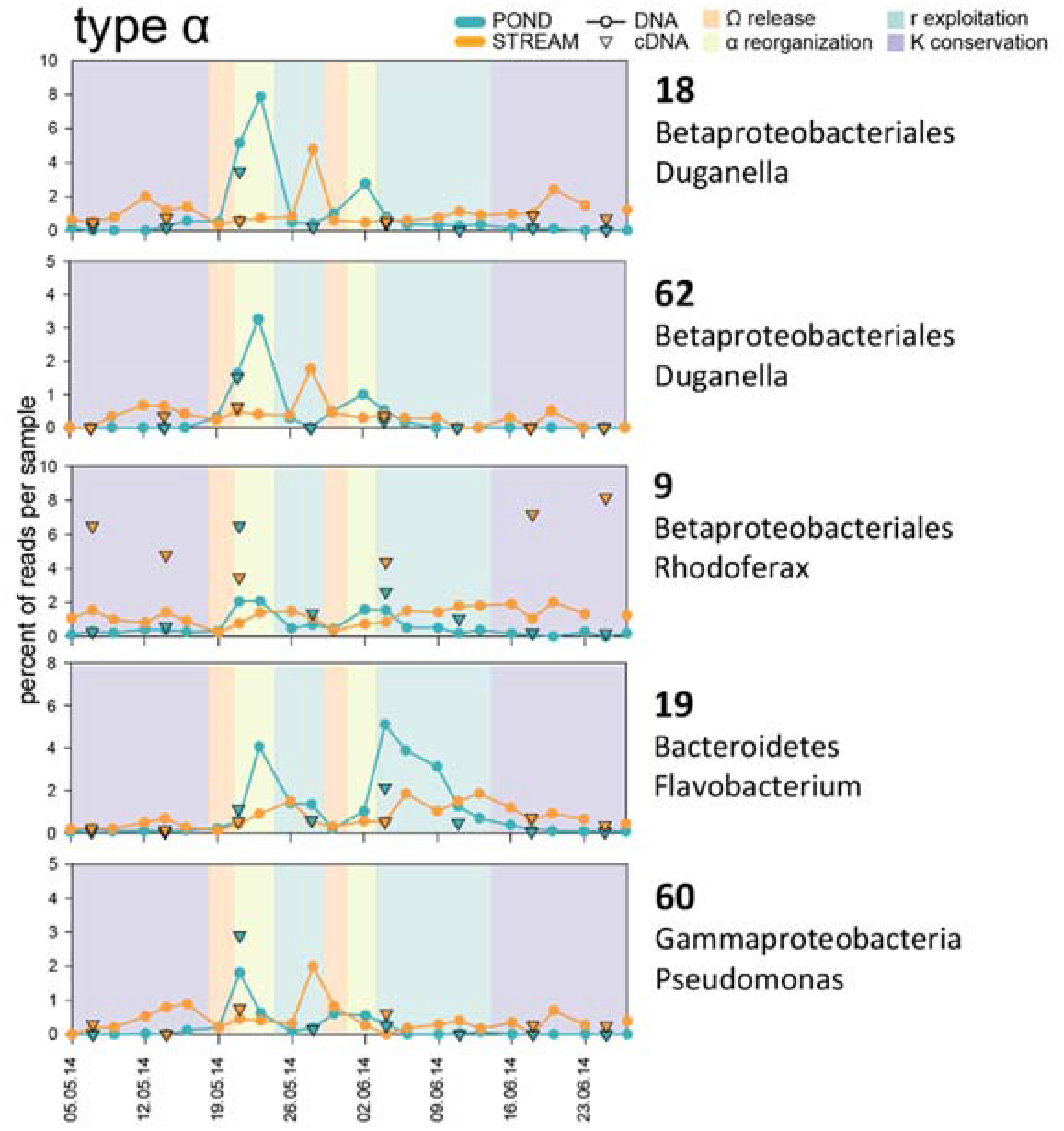
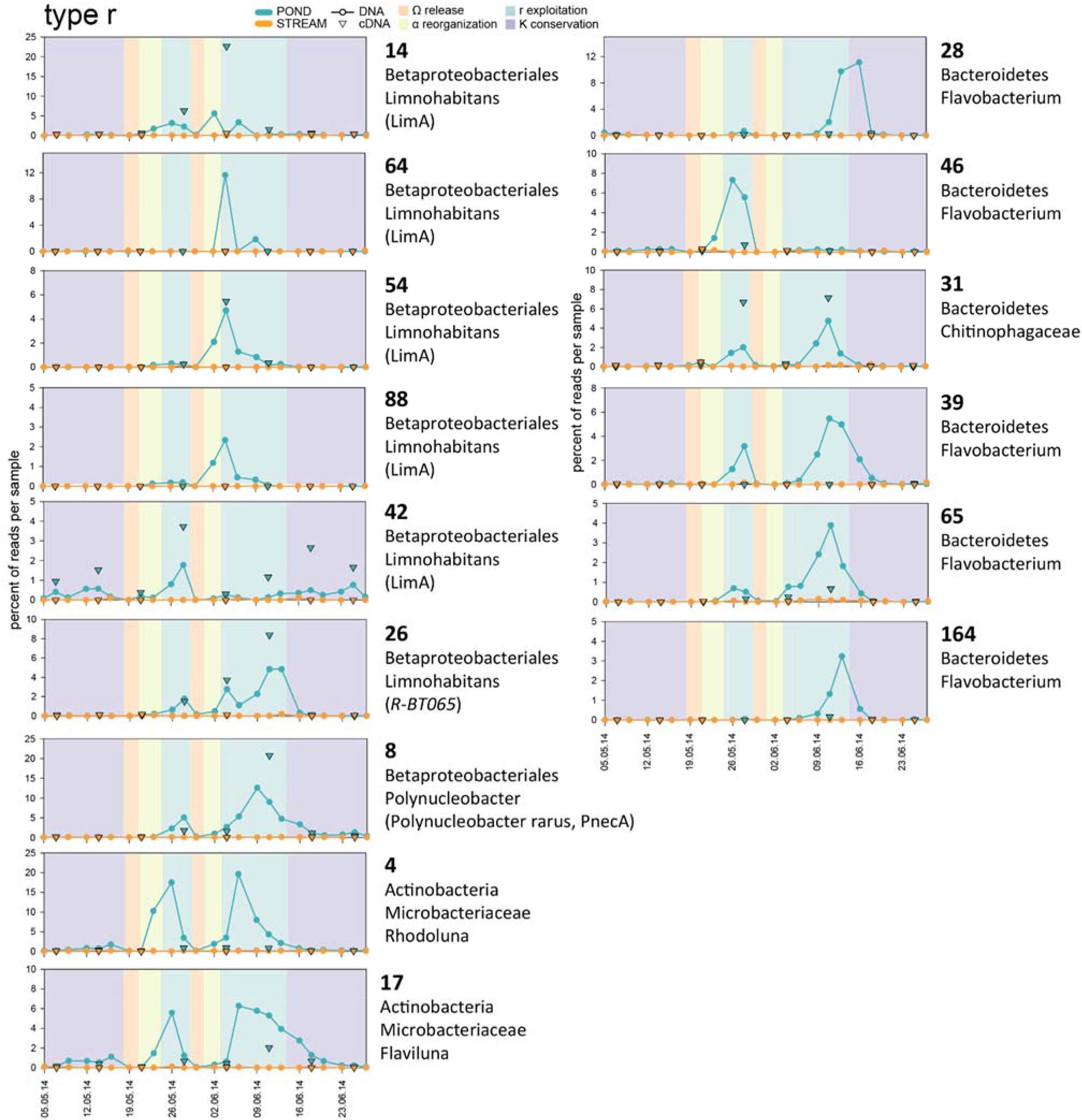
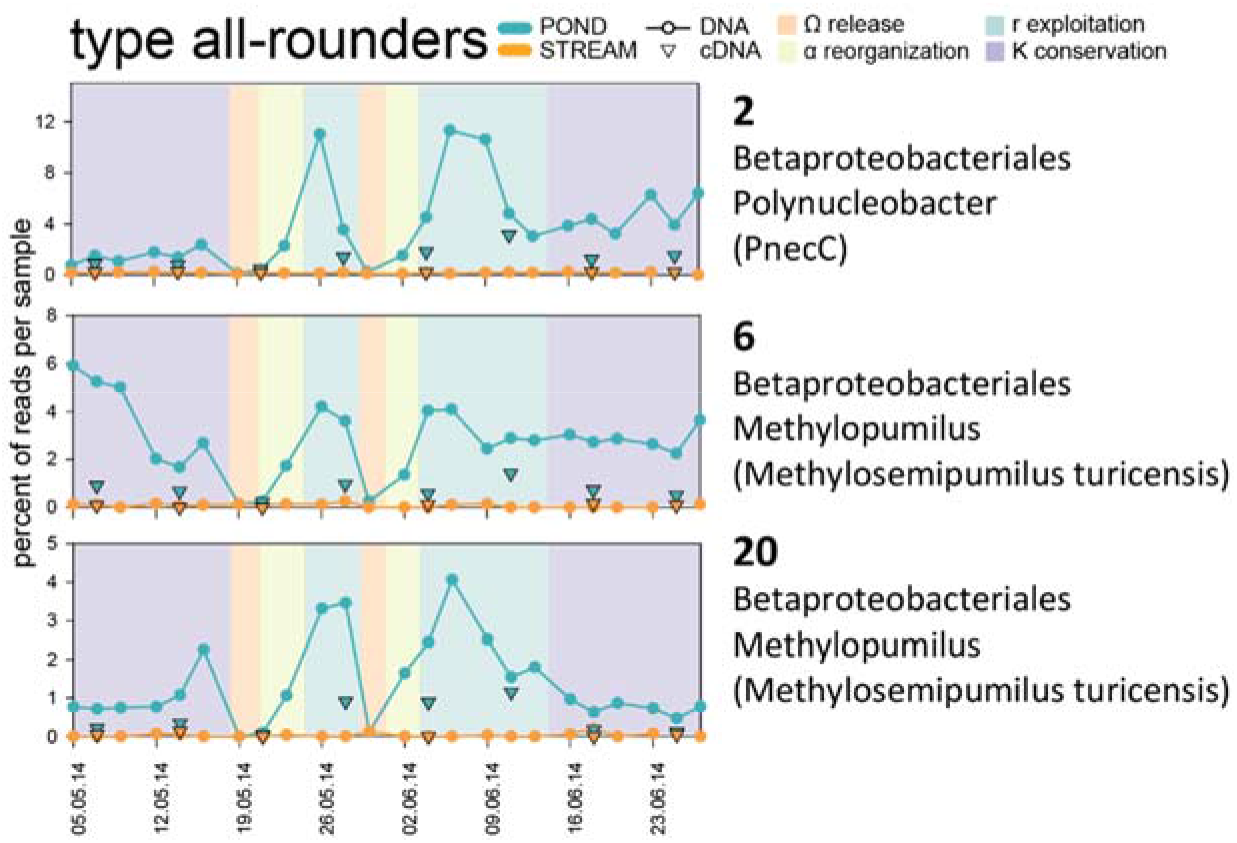
Different dynamic types of read proportions among pond ASVs with highest read abundances. Circles connected with lines represent percentages of 16S rDNA amplicon reads, triangles represent percentages of 16S rRNA amplicons reads. Samples from the pond are depicted in blue, samples from the stream in orange. Background colors indicate different phases of the succession: violet – ‘K’ conservation phase, orange – ‘Ω’ collapse and release phase, light green – ‘α’ reorganization phase, and aquamarine – ‘r’ exploitation phase. Displayed taxonomy is based on naive Bayesian classifier method and trained Silva SSU database release 132 incorporated in the DADA2 pipeline; additionally, if available, 100% identity matches obtained with BLAST are shown in brackets, and CARD-FISH probes matching the ASVs (RBT-065 and opitu-346) are indicated in italics. **a: type K**, ASVs detectable only during conservation phase, *K-*strategists. **b: type K(r)**, ASVs displaying highest read proportions in conservation phases and already detectable in exploitation phases (fast recovering *K-*strategists). **c: type Ω**, ASVs displaying highest read proportions during the EREs (emigrants) that do not belong to the most common pond ASVs. **d: type α**, ASVs displaying highest reads proportions during reorganization phase (inlet associated *r-*strategists). **e: type r**, ASVs displaying highest reads proportions during exploitation phase (pond associated *r-* strategists). **f: type all-rounders**, ASVs displaying comparable read proportions during all phases except EREs.

**Extended Data Fig. 6:**
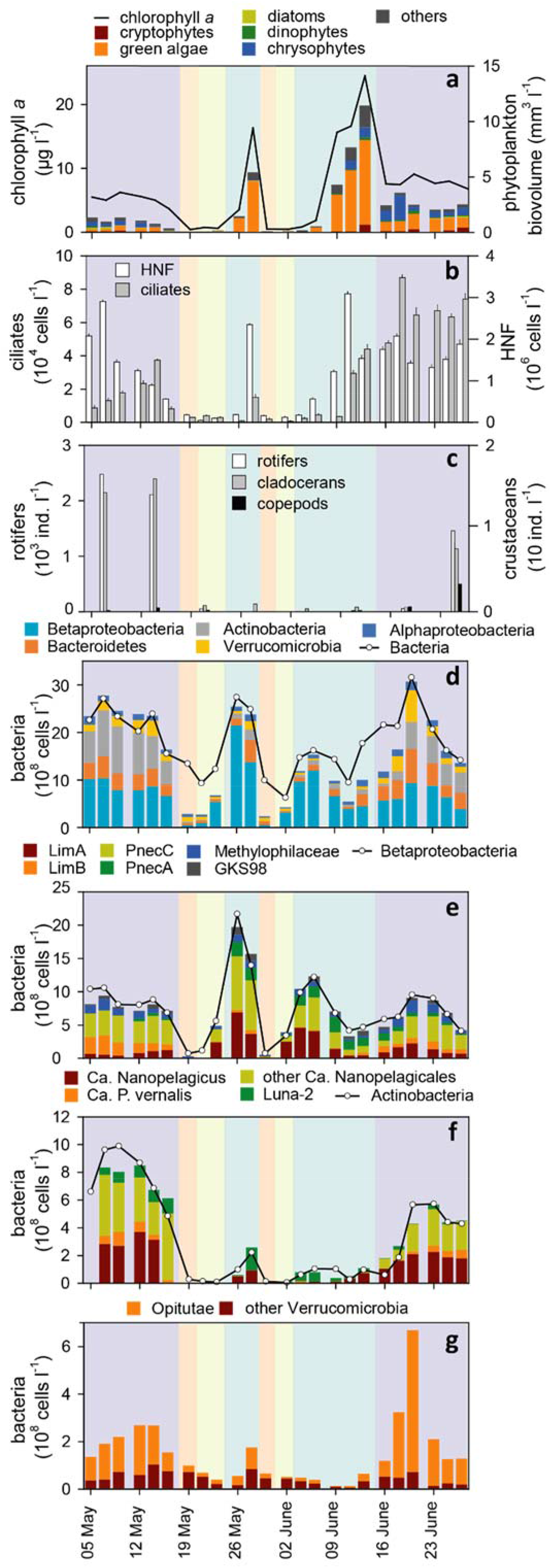
Aerophotographs of Jiřická Pond. **a:** Dam site; **b:** Stream site. Photographs were kindly provided by Petr Znachor.

**Extended Data Fig. 7:**
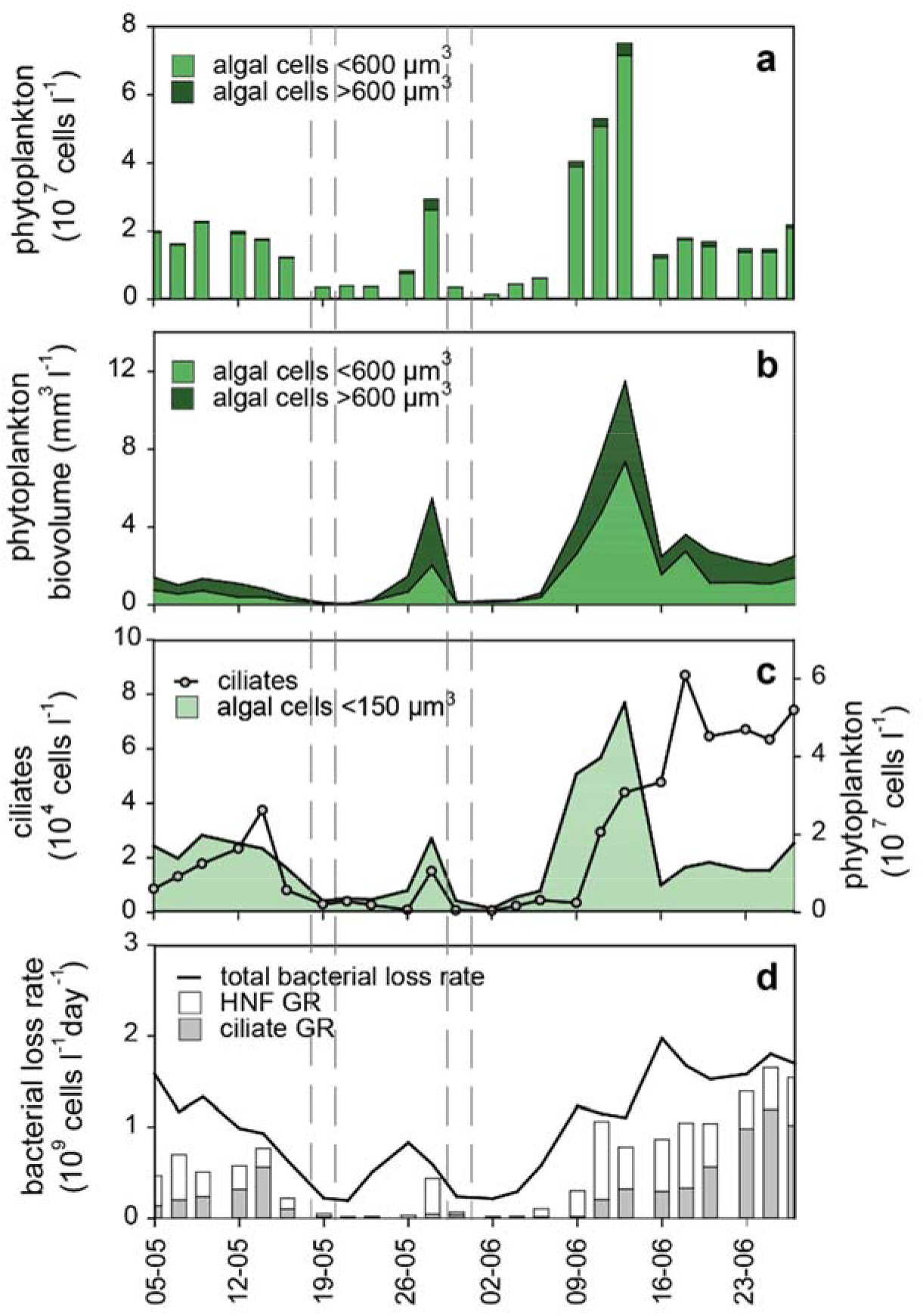
Changes in different size fractions of algae in the context of ciliate dynamics and contribution to the total bacterial loss. **a, b:** Abundances and biovolumes of algal cells <600 μm^3^ and >600 μm^3^, respectively (i.e., suitable or not suitable for ciliate grazing). **c:** Abundances of ciliates and the most dynamic fraction of algae <150 μm^3^ (represented mostly by small *Chlamydomonas* spp.). **d:** Total bacterial loss rate and HNF and ciliates grazing rates (GR). The dashed lines indicate the two EREs.

**Extended Data Fig. 8:**
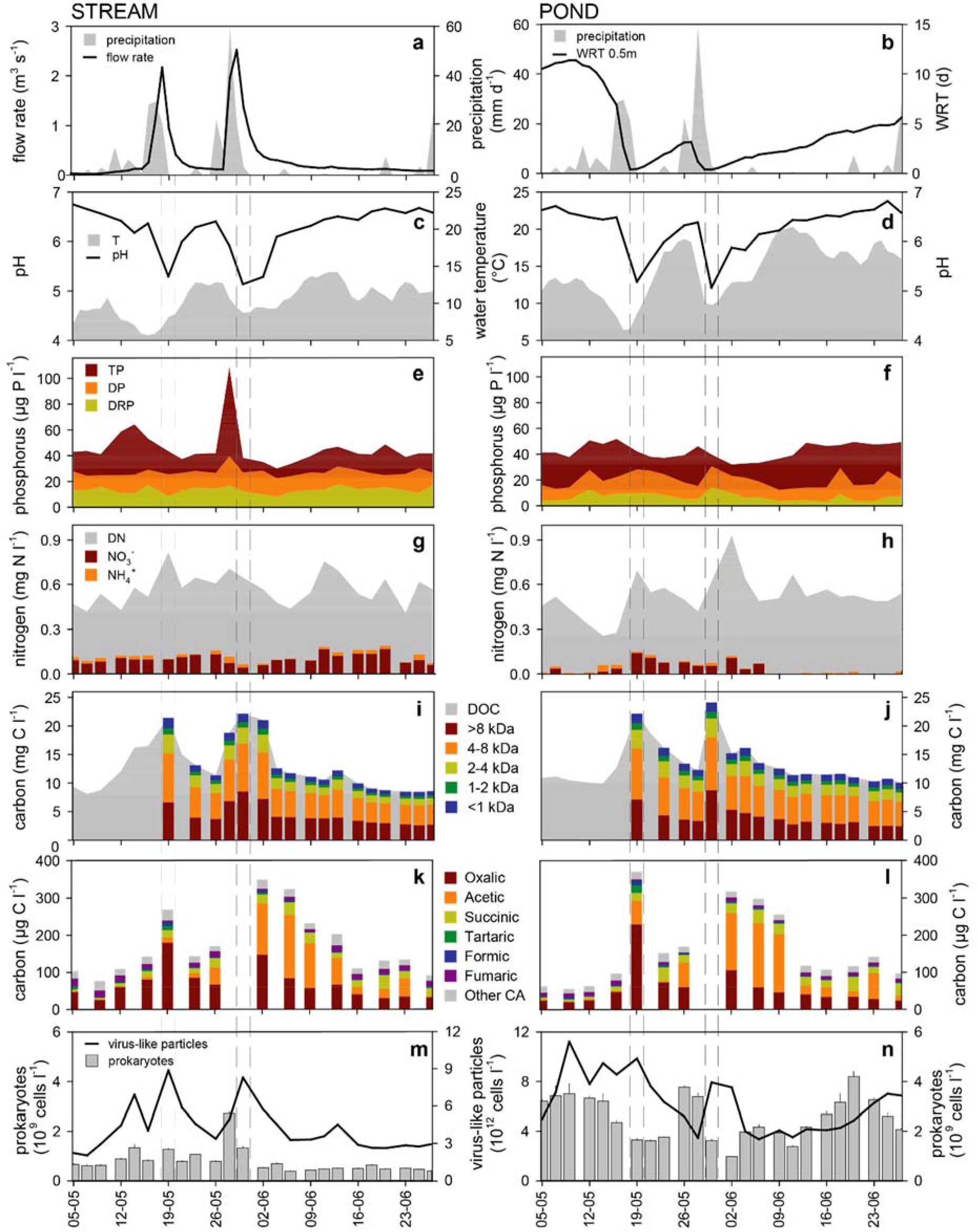
Hydrological, physicochemical and microbial parameters measured in the stream (left: **a, c, e, g, I, k, m**) and pond (right: **b, d, f, h, j, l, n**). Extended version of Figure 1. **a, b**: Precipitation (grey shade) in the watershed of Jiřícká Pond, stream flow rate (**a**) and water retention time (WRT) at 0.5m depth in the pond (**b**). **c**, **d**: Water temperature and pH. **e**, **f**: Concentrations of phosphorus fractions, TP – total phosphorus, DP – dissolved phosphorus, DRP – dissolved reactive phosphorus. **g**, **h**: Concentrations of nitrogen fractions, DN – dissolved nitrogen. **i**, **j**: Concentrations of dissolved organic carbon (DOC) and DOC size fractions. **k**, **l**: Concentrations of carboxylic acids (CA). **m**, **n**: Abundances of prokaryotic cells and virus-like particles. Dashed lines indicate two flood events during the sampling campaign.

**Extended Data Fig. 9:**
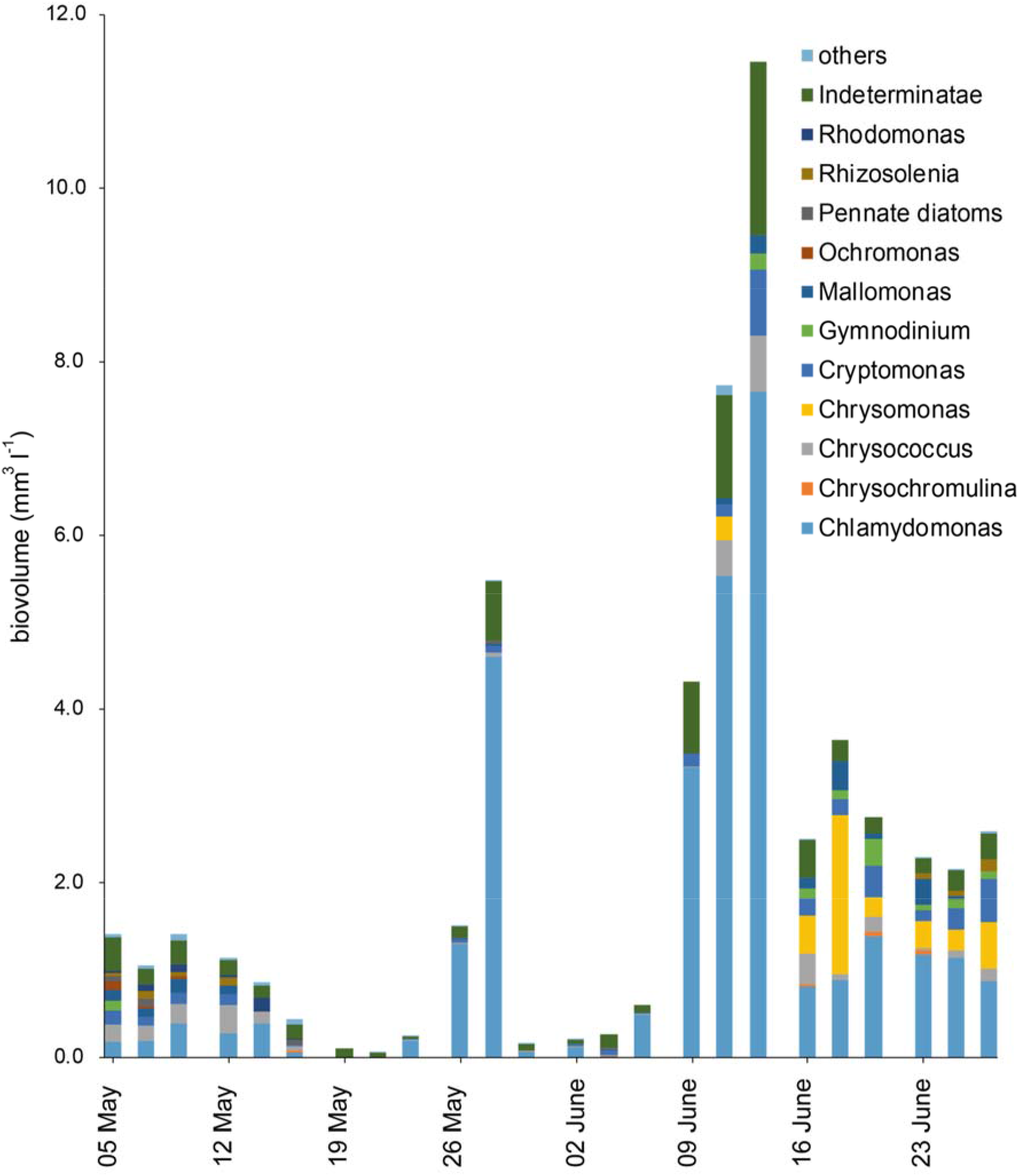
Composition of phytoplankton taxa (biovolume).

**Extended Data Fig. 10:**
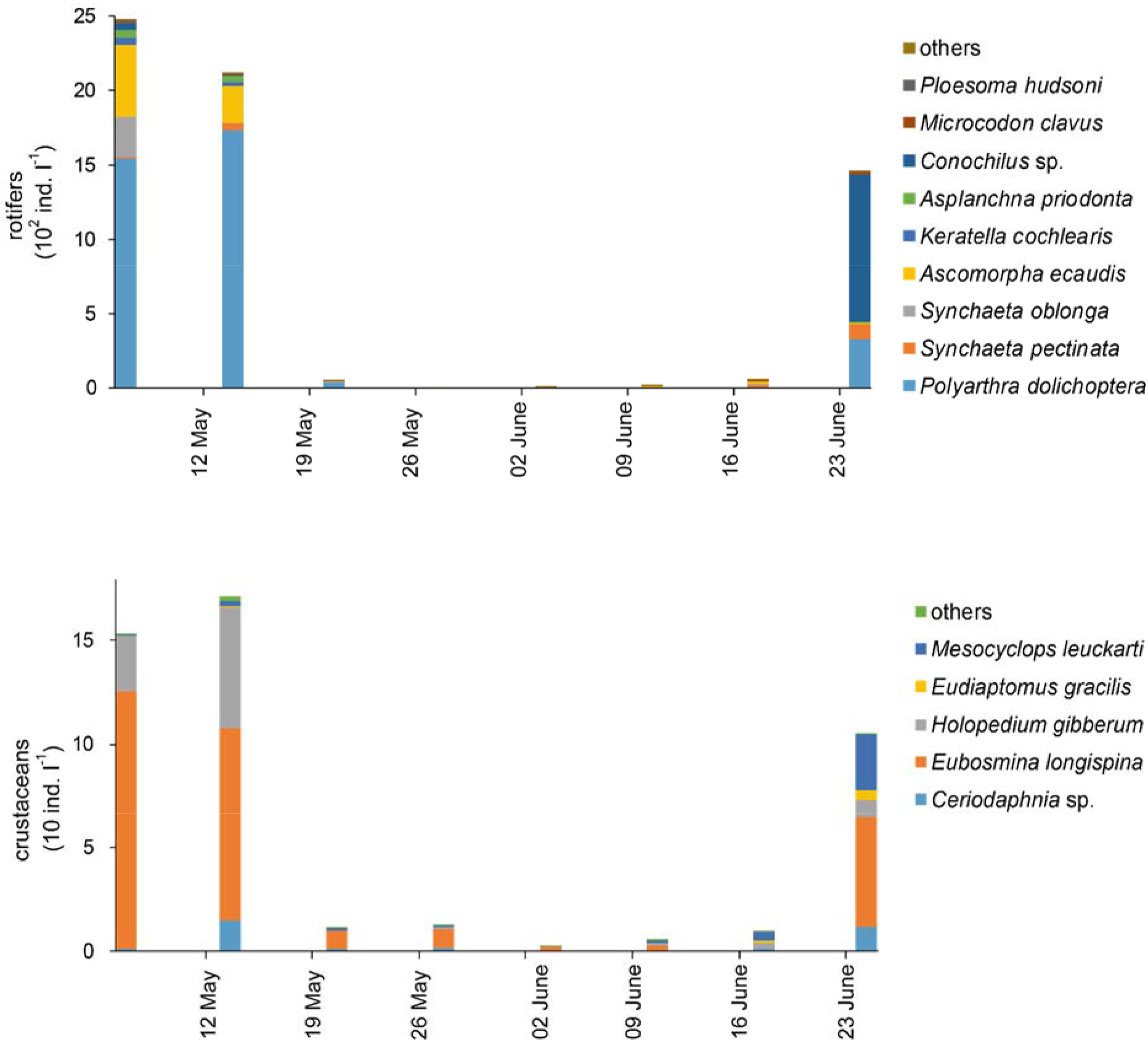
Composition of zooplankton taxa affiliated with rotifers and crustaceans.

**Extended Data Fig. 11:**
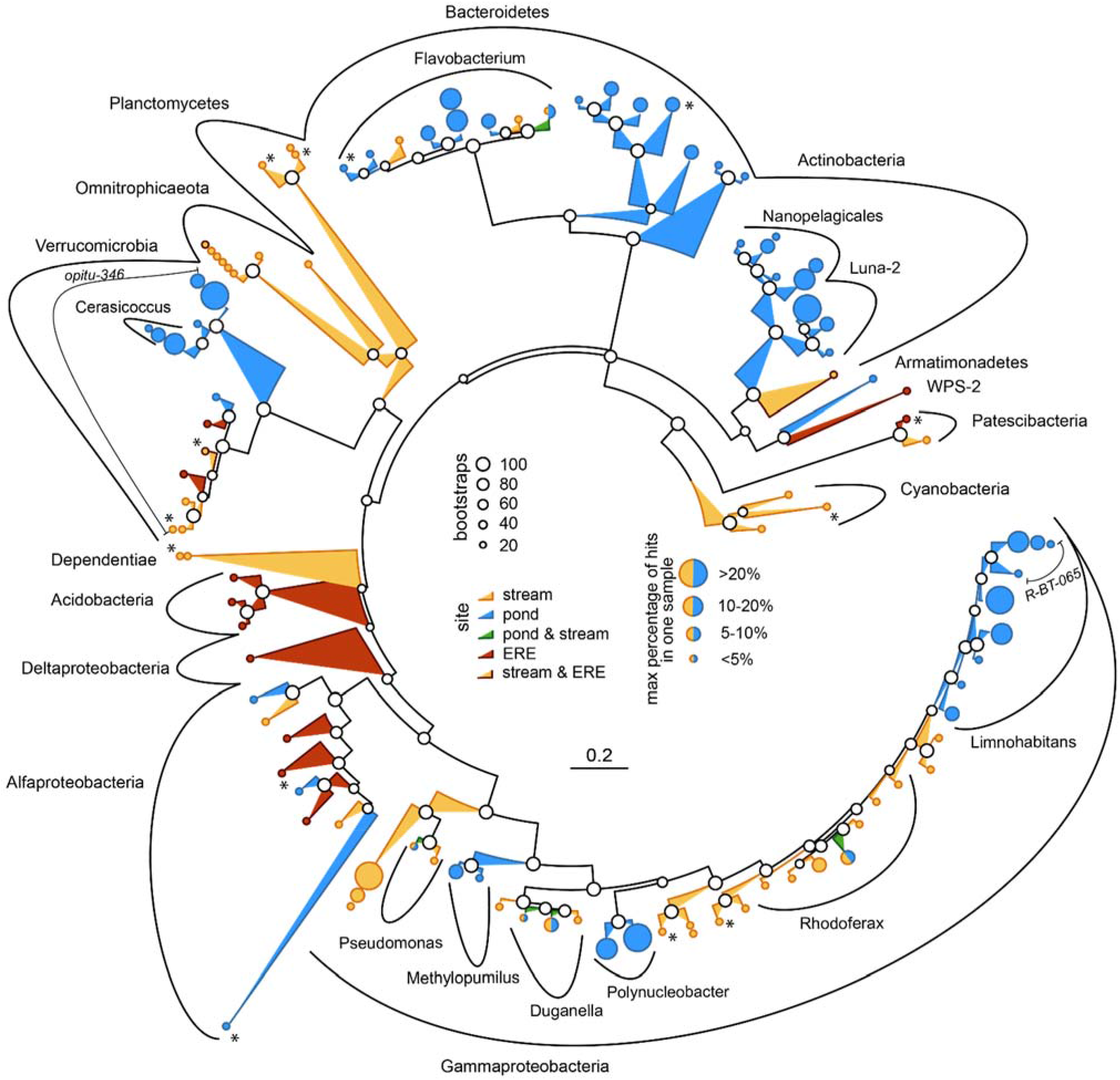
Bootstrapped maximum likelihood tree of 16S rRNA gene sequences of the closest relatives of the 50 most abundant ASVs detected in stream and pond samples and most abundant EREs genotypes. The colors of the branches and ASVs correspond to the source: orange – stream, blue – pond, green – pond and stream, red – EREs, orange with brown lines – stream and EREs. ASVs are depicted as circles with different diameters representing the maximal percentages of corresponding reads, ASVs shared between pond and stream are shown as divided circles. Asterisks indicate ASVs with low identity values to the closest relatives (90-97%). Specific CARD-FISH probes are shown in italics. Bootstraps (100 repetitions) are indicated by different sized circles at the nodes, the scale bar at the bottom applies to 20% sequence divergence.

**Extended Data Fig. 12:**
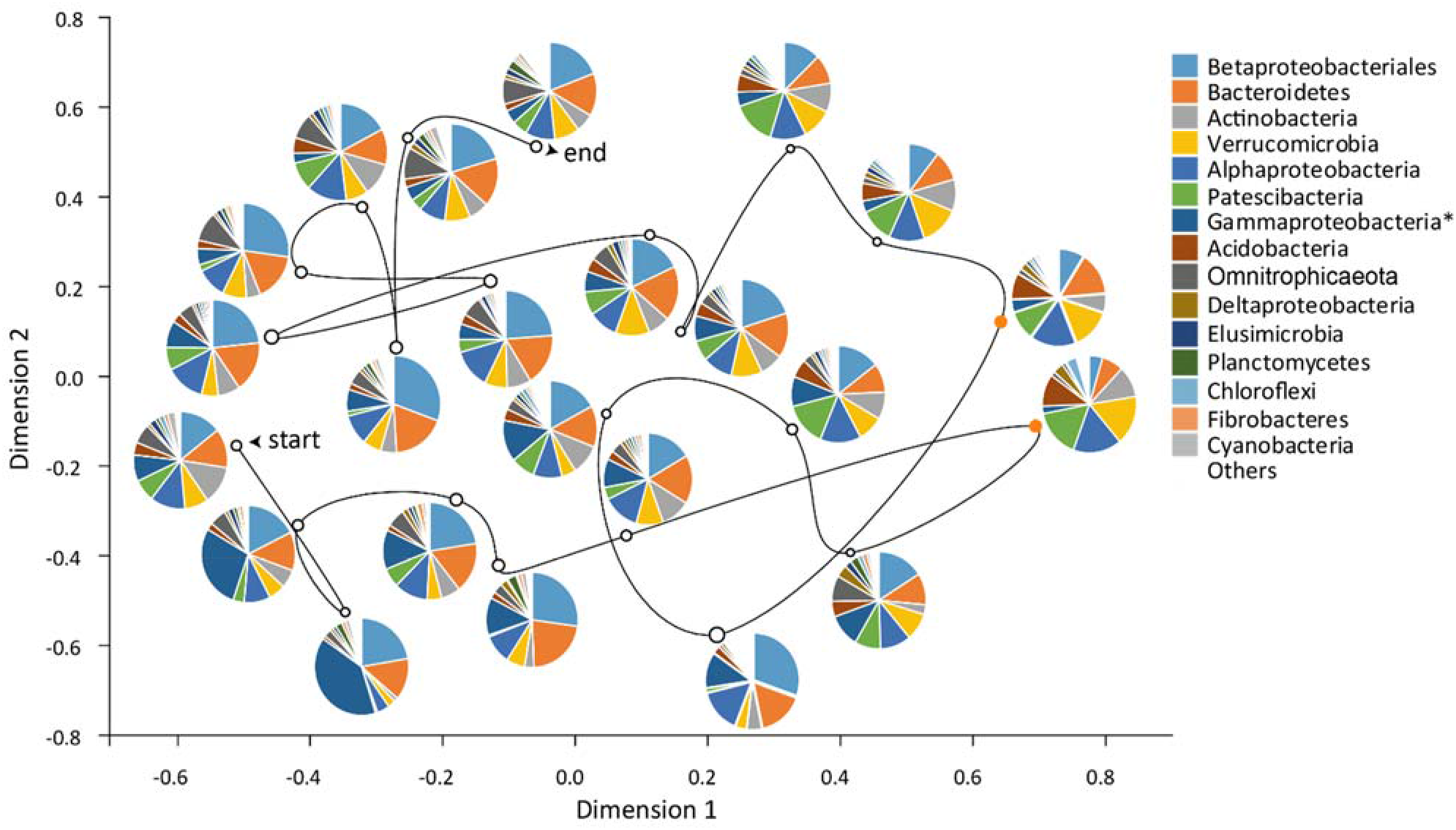
non-metric Multidimensional scaling (NMDS) based on Bray-Curtis distances of bacterial community compositions in the stream samples (16S rRNA gene amplicons), Kruskal's stress: 0.2; (**start**) and (**end**) indicate the first and last samples, respectively. The line connecting samples follows a chronological order. The diameters of the circles are proportional to the Pielou’s evenness of the samples (varying between 0.27 and 0.30). Orange circles indicate samples taken during the EREs. Pie charts depict the contribution of large taxonomical groups. *Gammaproteobacteria are shown without Betaproteobacteriales, which are presented as a separate group.

**Extended Data Fig. 1:**
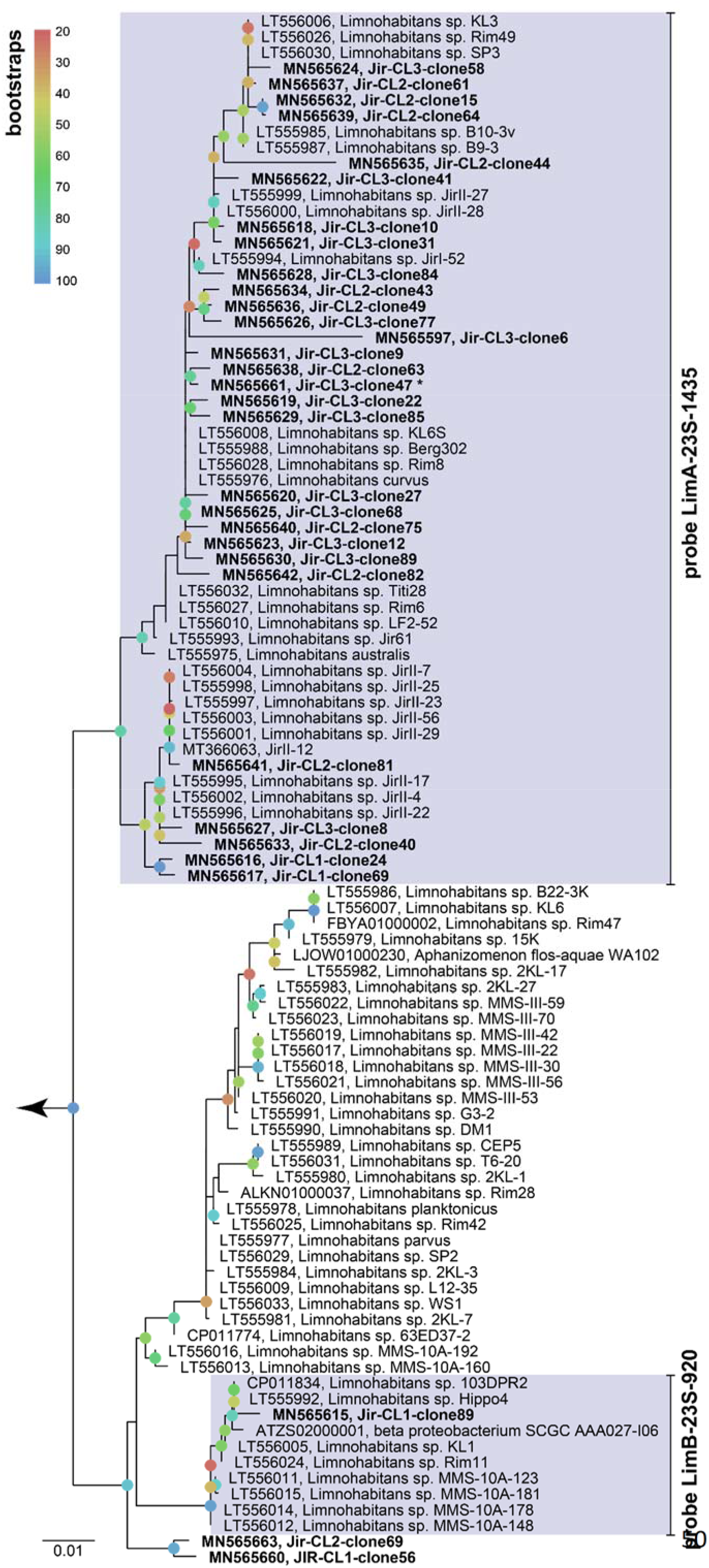
Bootstrapped maximum likelihood tree of 23S rRNA gene sequences of the genus *Limnohabitans* with marked target hits for probes LimA-23S-1435 (lineage LimA ^39^ and LimB-23S-920 (lineage LimB; this study). Sequences obtained from clone libraries are depicted in bold (CL1: 12.05.2014; CL2: 26.05.2014 and CL3: 02.06.2014). Bootstraps (100 repetitions) are indicated in different colors, branches with bootstraps <20% were multifurcated. The scale bar at the bottom applies to 1% sequence divergence.

**Extended Data Fig. 14:**
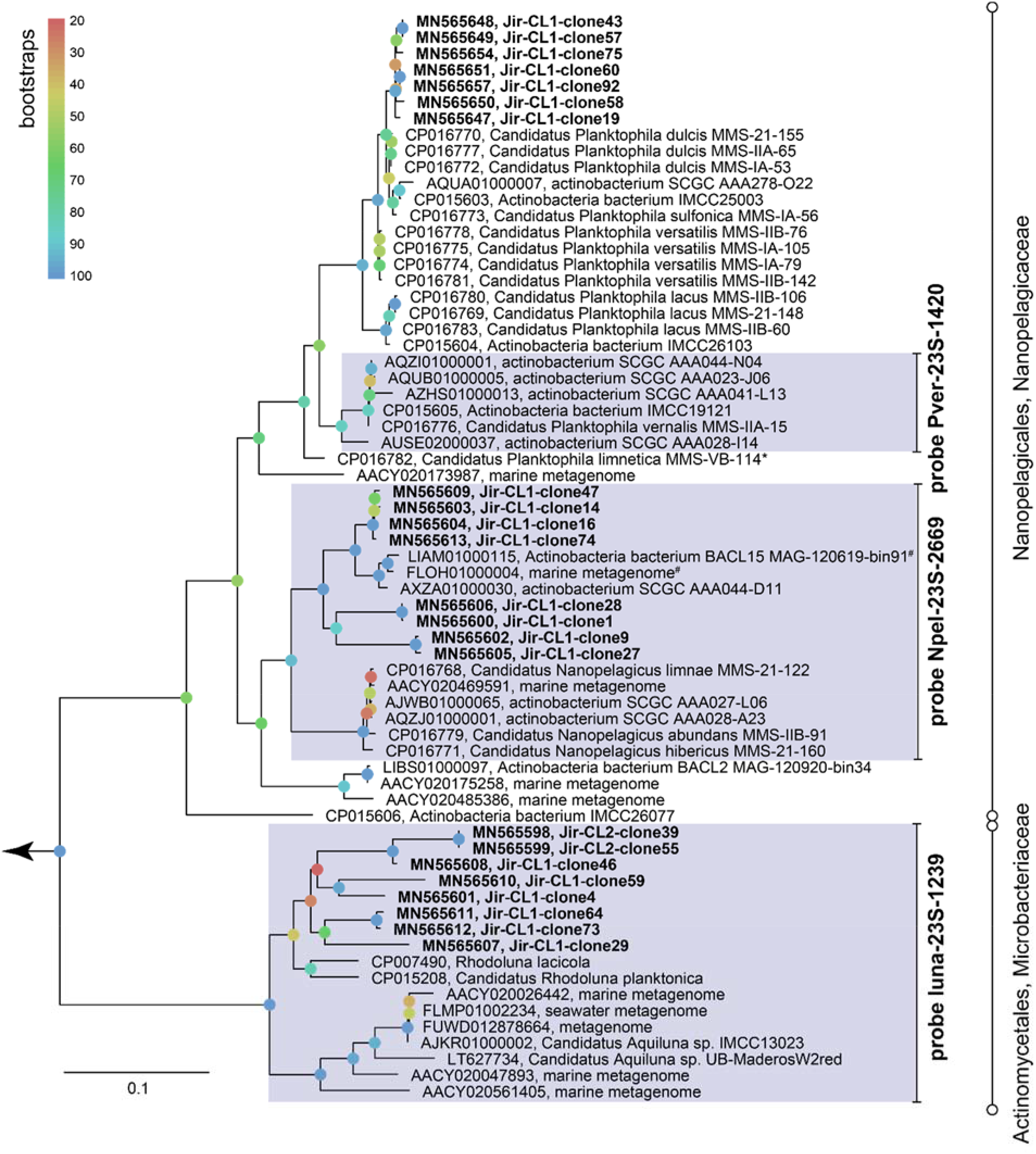
Bootstrapped maximum likelihood tree of 23S rRNA gene sequences affiliated with Actinobacteria with marked target hits for probes Pver-23S-1420 (‘*Ca*. *Planktophila vernalis*’^27^), Npel-23S-2669 (‘*Ca*. Nanopelagicus’^27^), and luna-23S-1239 (luna2 lineage of Microbacteriaceae; this study). Sequences obtained from clone libraries are depicted in bold (CL1: 12.05.2014; CL2: 26.05.2014 and CL3: 02.06.2014). Bootstraps (100 repetitions) are indicated in different colors; the scale bar at the bottom applies to 10% sequence divergence.

**Extended Data Fig. 15:**
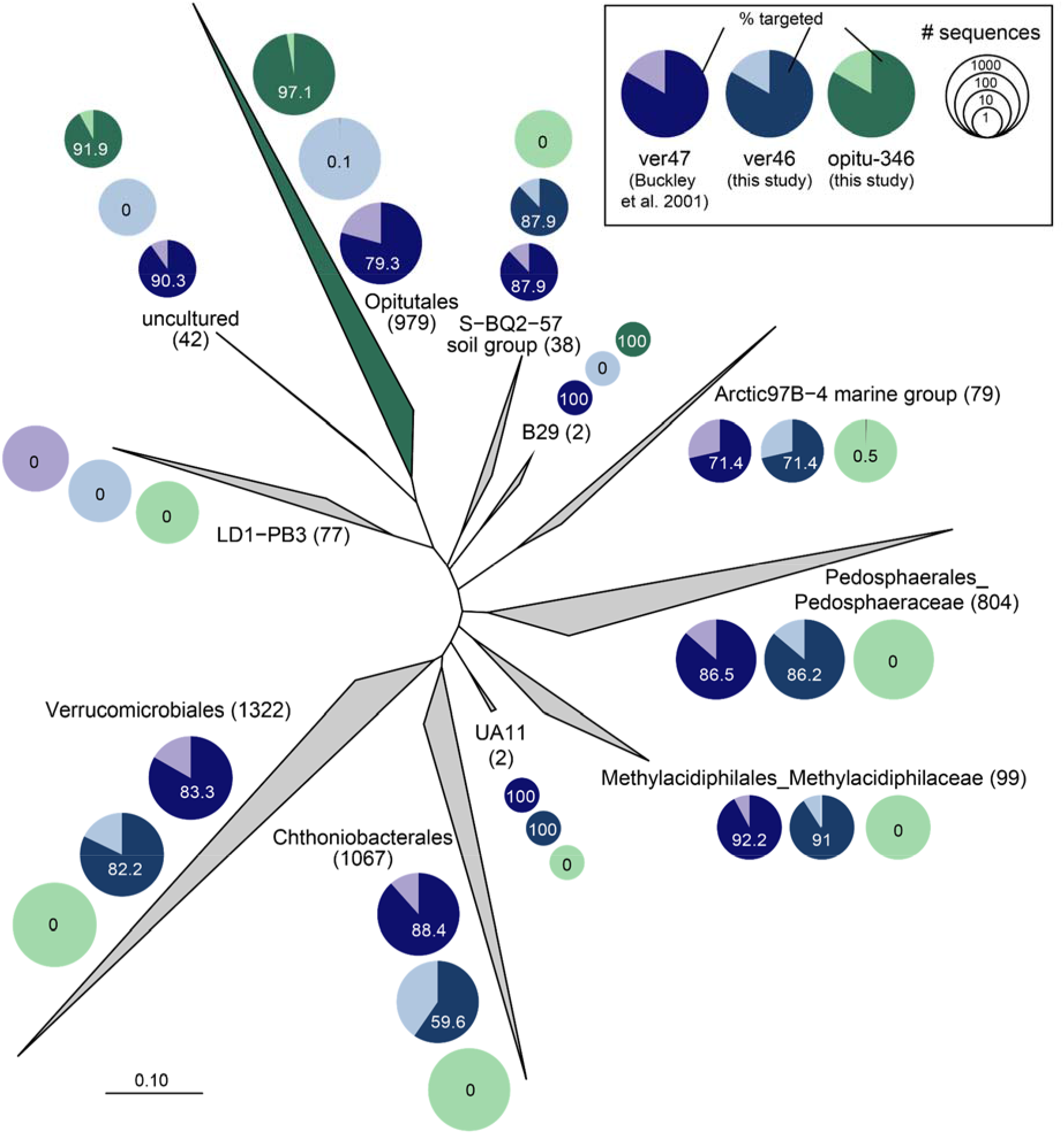
Extract of the maximum parsimony guide tree of SILVA_ 132_SSURef_NR99 depicting the coverage of specific CARD-FISH probes for different lineages of Verrucomicrobia. The number of sequences within each collapsed branch is indicated in brackets and by different sized circles. Coverage of general probe ver47^70^ is shown in turquoise, the modified version of this probe (ver46; this study) in blue, and probe opitu-346 (Opitutales, this study) in green. The scale bar at the bottom applies to 10% sequence divergence.

**Extended Data Fig. 16:**
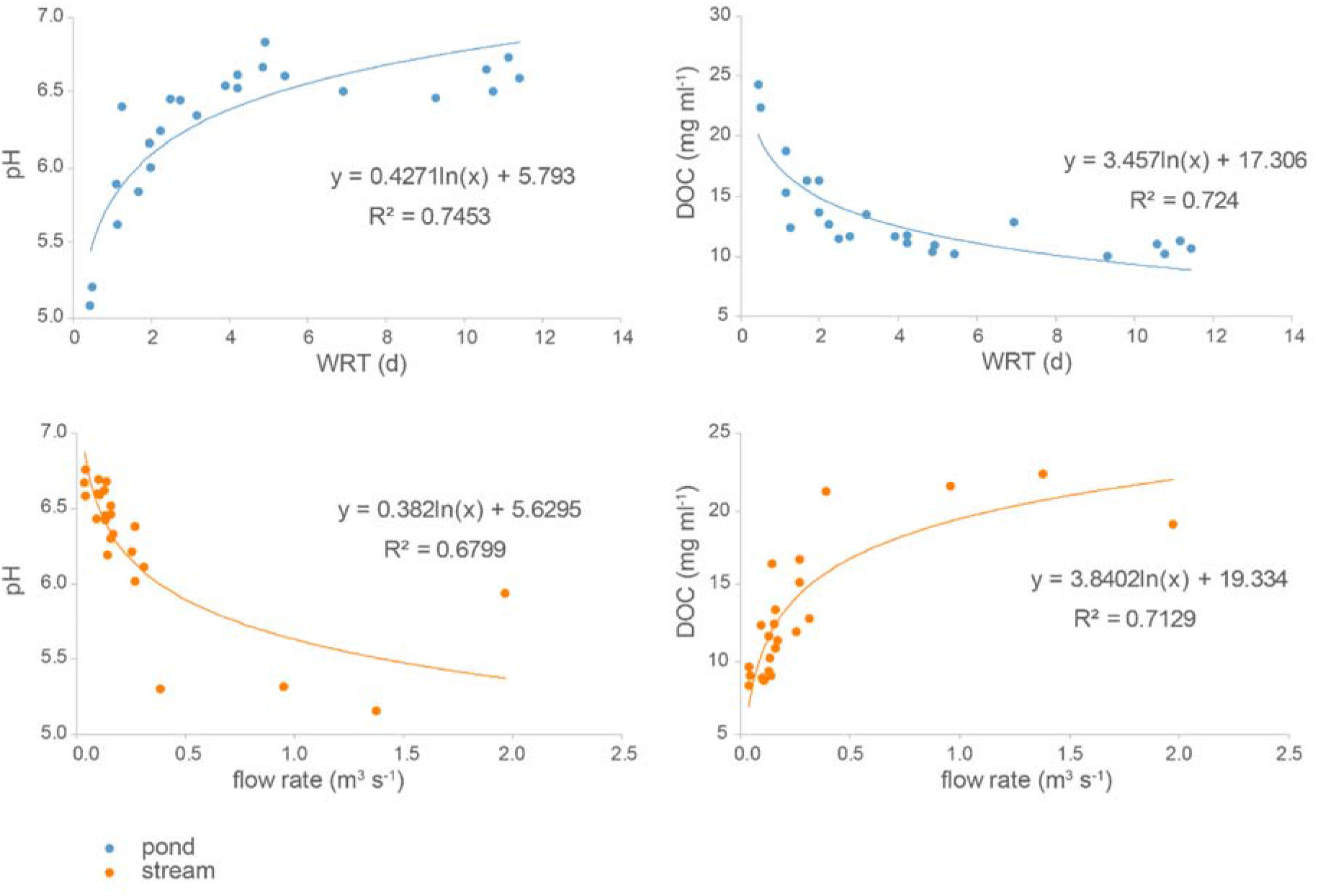
Relationship between hydrological and chemical parameters in pond and stream. Pond and stream data are depicted in blue and orange, respectively.

## Extended Data Text 1

### Sampling site

Humic pond Jiřická (48.6189N, 14.6731E; 892 m a.s.l.) is a small water body in Novohradské Mountains (CZ) with a surface area of 0.036 km^2^ and an average WRT of 9 days^23, 24^. The pond is located in the headwaters of Malše river and belongs to the catchment of Vltava river (tributary of Elbe), where an increasing frequency of flood events was observed in the last 40 years^13^. Jiřická represents a typical headwater body of Hercynian Mountains of Central Europe and can serve as a model ecosystem to study flood-related ecological processes. Two locations were intensively sampled (three times a week) during precipitation driven, hydrologically very dynamic period between May 5^th^ and June 27^th^ 2014 to access flood-associated changes in the plankton community: i) station pond, an epilimnetic station, located approx. 50 m from the dam at the deepest part of the pond, and ii) station stream, the main riverine tributary sampled 500 m upstream from the pond.

### Chemistry

Most physicochemical characteristics in both stream and pond were related to hydrological and meteorological parameters (Extended Data Fig. 3 and 8). For example, total concentrations of phosphorus detected in the stream responded to the precipitation levels, while concentrations of inorganic nitrogen species in the pond followed the hydrological dynamics (Extended Data Fig. 3 and 8). Moreover, we observed a logarithmic relationship between hydrology and dissolved organic carbon (DOC) concentrations as well as pH at both sites (Extended Data Fig. 16). Amounts of chromophoric dissolved organic matter (CDOM) and DOC, showing overall correlations in the pond (r^2^=0.97), increased by a factor of 1.7 (ERE1) and 2.0 (ERE2) compared to pre-flood values. High molecular weight fraction of DOC (> 8 kDA) also increased in percentages during and shortly after EREs. At the same time, the composition of carboxylic acids (forming a substantial part of the smallest DOC fraction), especially the proportions of acetic and oxalic acids, varied substantially between different EREs (Extended Data Fig. 8l). Comparison of main physicochemical parameters between pond and inlet revealed only small numbers of significantly distinctive characteristics, namely temperature and nutrients concentrations. The soluble inorganic fractions of nitrogen and phosphorus were significantly lower in the pond than in the inlet showing high consumption rates of the nutrients in the pond, especially during stable hydrological conditions. At the same time, we did not find significant differences between pond and inlet in the concentrations of TP and DOC, and calculated budgets for them were rather neutral. Similarly, the concentrations of entire dissolved nitrogen (DN) in the pond and inlet did not differ substantially.

### Extended description of succession phases

#### Conservation phase (K)

Dominant taxa were affiliated with typical lake microbes including Actinobacteria, Verrucomicrobia, Bacteroidetes and Betaproteobacteria (Extended Data Fig. 3a-b), e.g. ‘*Ca.* Planktophila’ spp. (amplicon sequence variants (ASVs) 3, 12, 15, 21, 38 and 53), Opitutae (ASVs 1, 8, 112, 3170, 3602), *Limnohabitans* lineage LimB (ASV 30), *Polynucleobacter* lineage PnecC (ASV 2). The majority of sequences (65.5%) detected in this phase had closest relatives reported from aquatic environments, whereof 31% originated from the lake water column. Chlorophyll *a* concentrations and total phytoplankton biovolumes were relatively low and ranged from 3.82 to 9.23 μg l^−1^ and from 0.44 to 3.65 mm^3^ l^−1^ in first and second K phases respectively (Extended Data Fig. 6a, 7b). The start of the second K phase was associated with a collapse of phytoplankton visible by a drop in biovolume from 11.5 to 2.5 mm^3^ l^−1^, mainly due to a decrease of *Chlamydomonas* spp. abundance (from 54.6∙10^6^ to 3.8∙10^6^ cells l^−1^, Extended Data Fig. 5c). Protistan and zooplankton dynamics differed between both conservation phases: ciliates were present in low numbers before ERE1 (<3.7∙10^4^ ciliates l^−1^) and reached high abundances in the second K phase (4.8∙10^4^ to 8.7∙10^4^ l^−1^), while the opposite was found for zooplankton (Extended Data Fig. 6b-c). During the second conservation stage, ciliate community composition shifted from consumers of flagellated algae and protozoa (i.e. prostomatids feeding on *Chlamydomonas* spp. and partially HNF) to bacterivorous and omnivorous ones. Thus, the bacterivory rate of ciliates was 3∙10^8^ bacterial cells d^−1^ l^−1^ at the ciliate abundance maximum (on June 18^th^), and four times higher (1.2∙10^8^ bacterial cells d^−1^ l^−1^) one week later at lower ciliate densities (Extended Data Fig. 7c-d). Generally, bacterial disappearance rates mainly due to microzooplankton grazing were higher in the second stable phase (ranging between 0.635∙10^9^ and 1.80∙10^9^ bacterial cells d^−1^ l^−1^ Extended Data Fig. 7d).

#### Mass effects driven collapse phase (Ω)

The samples taken during the EREs in the pond were characterized by very low prokaryotic densities and high diversity (Fig. 1, Extended Data Fig. 4). This was reflected in the presence of many bacterial ASVs with low read percentages, i.e. within Verrucomicrobia, Alphaproteobacteria, and taxa previously almost absent in the pond, i.e. Acidobacteria, Deltaproteobacteria, Patescibacteria, WPS-2 and Omnitrophicaeota (Extended Data Fig. 5c, 11). The most abundant (% of reads) ASVs were represented by the Solirubrobacteraceae family of Actinobacteria, Acidobacteria, Opitutaceae, Saccharimonadales and WPS-2. Most of these phylotypes were not present in the stream at low flow conditions and thus might represent emigration from habitats connected to the stream only during EREs. However, ASVs 102, 212 and 121 were present in the stream at all hydrological regimes and seemed to be active as indicated by decent numbers of cDNA reads.

#### Reorganization phase (α)

This phase was characterized by low WRT in the pond (<2.0 d). Fast-growing bacterial phylotypes (*r*-strategists) persistent in the stream (Extended Data Fig. 5d) were dominating the pond microbial community. At the same time an increase in reads percentages of pond-specific *r*-strategists (Extended Data Fig. 5e) was observed, consisting of different ASVs affiliated with *Limnohabitans* lineage LimA, *Polynucleobacter rarus*, diverse Bacteroidetes, and Actinobacteria of the luna-2 lineage. While the majority of these ASVs had maxima after both EREs, several were growing to high densities only after ERE1 (ASVs 42, 46) or ERE2 (ASVs 28, 64, 54, 88, 164). Composition of carboxylic acids (CAs) also showed high heterogeneity during this phase: while oxalic and tartaric acid concentrations increased after both floods, elevation of acetic acid concentrations was observed only after ERE2 (Extended Data Fig. 3). This could be connected to alternations in pond specific *r-*strategists after EREs (e.g. ASVs associated with LimA and *Flavobacterium* spp., Extended Data Fig. 5e).

#### Exploitation-phase (r)

This stage was characterized by a shift from ASVs shared with the stream to exclusively pond specific ones (Extended Data Fig. 5d-e), e.g. within *Flavobacterium* species, or from stream-specific *Rhodoferax* spp. to pond-specific *Limnohabitans* spp.. Moreover, we observed fast recovery of two Microbacteriaceae ASVs typical for freshwaters (*Rhodoluna* and *Flaviluna*^78^), which was also reflected in elevated CARD-FISH abundances of luna-2 lineage (Fig. 2, Extended Data Fig. 6f). This lineage of Actinobacteria displayed divergent patterns with peaks in r and K phases associated with different ASVs (Extended Data Fig. 5b, e). Although bacterial densities reached pre-flood abundances during the first r phase (Fig. 1), they did not raise above 2.2∙10^9^ l^−1^ during the second one and the minimum in bacterial abundances on June 11^th^ (1.4∙10^9^ l^−1^) corresponded well to a maximum of bacterivorous HNFs (3.1∙10^6^ l^−1^; Extended Data Fig. 5b, d). Phytoplankton displayed exponential growth during exploitation-phases and reached their maxima on May 28^th^ and June 13^th^, respectively (Extended Data Fig. 6–8). Chemical and hydrological characteristics were comparable with stable conditions and removal of soluble inorganic nutrients fractions recovered to pre-disturbance level (Fig. 4, Extended Data Fig. 3 and 8).

### Zoo- and phytoplankton dynamics

Zooplankton was mainly represented by *Polyarthra dolichoptera* (rotifers) and *Eubosmina longispina* (crustaceans), which are common members of shallow lake plankton^79^ (Extended Data Fig. 6c, 9). Maxima of these species were often observed prior dominance of *Daphnia* spp. in spring successions^15^, corresponding well to their maxima at the beginning of our study. However, the reassembly of zooplankton observed after the second ERE resulted in an alternation of composition characterized by high densities of *Conochilus unicornis* and establishment of a population of the omnivorous copepod *Mesocyclops leuckarti* (Extended Data Fig. 6c, 9) typical for late spring vs early summer^23^. *Conochilus unicornis*, a colony-forming rotifer, possesses high clearance rates of phytoplankton with a preferred size spectrum of 5-9 μm and seems to play an important role in the phytoplankton decimation after ERE2. At the same time, it is more resistant to an ingestion by *Mesocyclops leuckarti*, which likely control the population of *P. dolichoptera* in summer (adult females of *M. leuckarti* can consume more than 40 individuals of *P. dolichoptera* per day^80^).

During the whole study period, phytoplankton was dominated mostly by small flagellated species, which typically occur at the beginning of the vegetation season, forming a vernal peak of biomass in response to increasing light availability^43^. In our study, both EREs strongly modified the seasonal succession of phytoplankton, resulting in pronounced peaks dominated mostly by *Chlamydomonas* spp. one week after each ERE (Extended Data Fig. 5a, 8). A similar pattern was observed in response to two summer floods on reservoir phytoplankton^81^, where the authors reported that the first ERE virtually initiated the development of summer phytoplankton assemblage while the second one a month later reversed phytoplankton succession to an earlier stage. In Jiřická Pond, *Chlamydomonas* spp. abundance decreased one week after ERE2 (Extended Data Fig. 5a, 8) and later on, several Chrysophyte and Cryptophyte species co-existed.

### Stream bacteria

The stream planktonic microbial community differed significantly in structure and composition from the pond epilimnion (Fig. 2b, Extended Data Fig. 2, 12). Diversity indices observed in stream bacterial assemblages were substantially higher than those in the pond, except for flood situations (Extended Data Fig. 3). Low bacterial densities along with relatively high particle contents and a high number of bacterial groups, even at high taxonomical levels (Extended Data Fig. 12), rendered the CARD-FISH analysis in the stream very difficult and unreliable. Algal and zooplankton composition were not assessed in the stream because of the negligible density of these groups, e.g. chlorophyll-*a* concentrations in the stream never exceeded 2.34 μg l^−1^, that was 2-10 times lower than in the pond during both r and K phases.

## Extended Data Tables

**Extended Data Table 1.**
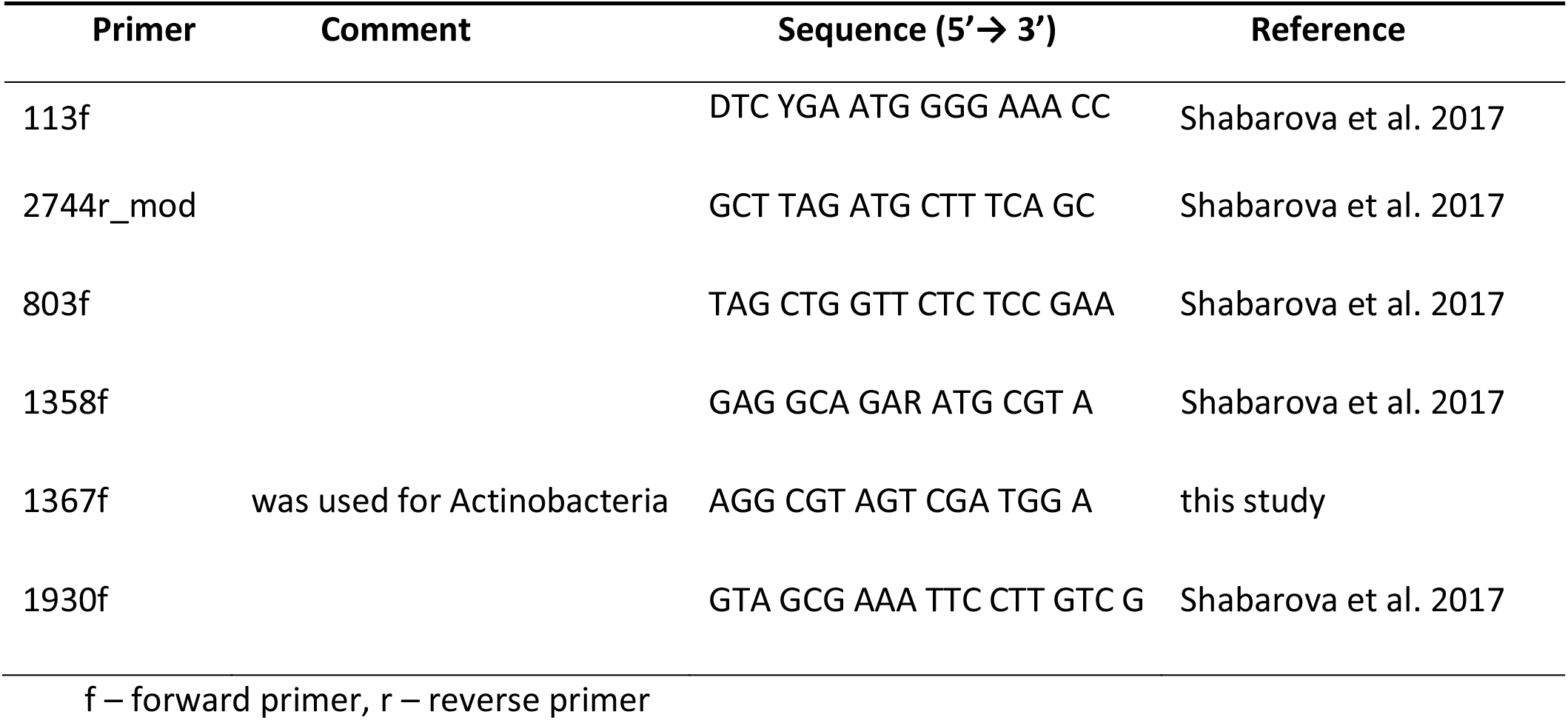
Primers used for amplification and sequencing of 23S rRNA genes

**Extended Data Table 2.**
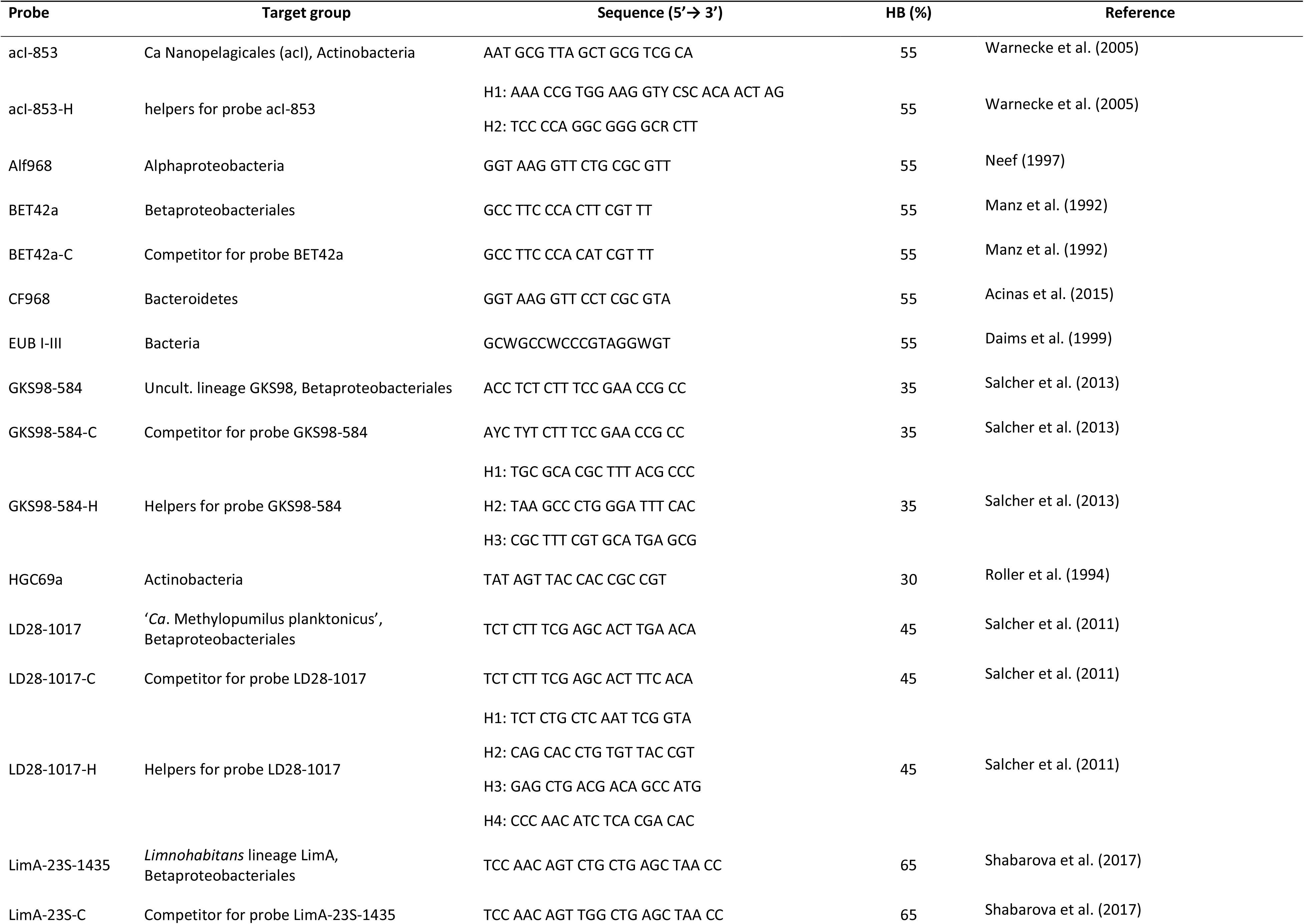
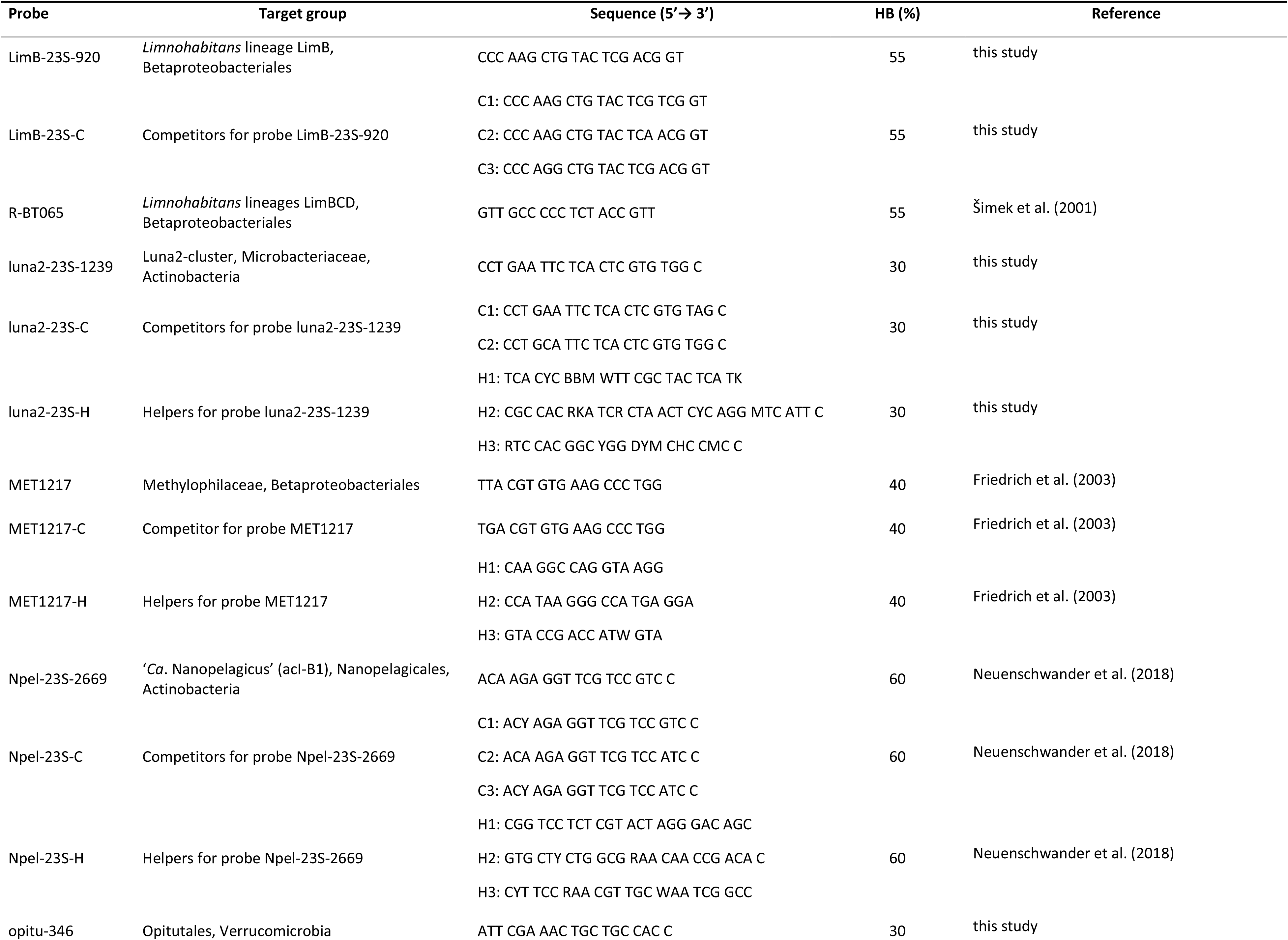
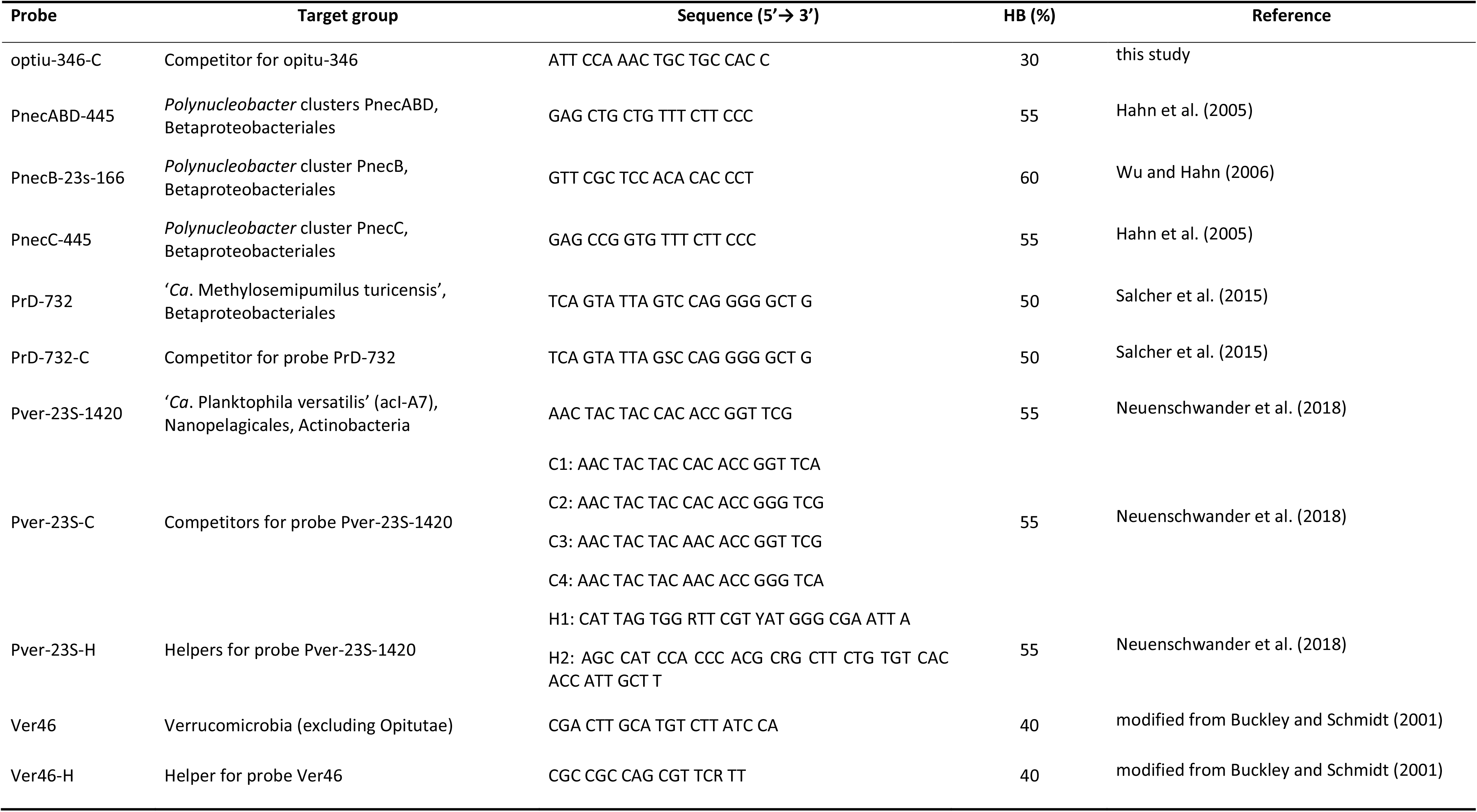
Probes, helpers and competitors used in this study. HB (%) – Formamide concentration in the hybridization buffer.

